# Sulfite oxidase deficiency causes persulfidation loss and H_2_S release

**DOI:** 10.1101/2024.03.13.584820

**Authors:** Chun-Yu Fu, Joshua B. Kohl, Filip Liebsch, Davide D’Andrea, Max Mai, Anna T. Mellis, Emilia Kouroussis, Tamás Ditrói, José Angel Santamaria-Araujo, Sin Yuin Yeo, Heike Endepols, Michaela Křížková, Viktor Kozich, Uladzimir Barayeu, Takaaki Akaike, Julia B. Hennermann, Peter Nagy, Milos Filipovic, Guenter Schwarz

## Abstract

Sulfite oxidase (SOX) deficiency is a rare inborn error of cysteine metabolism resulting in severe neurological damage. In patients, sulfite accumulates to toxic levels causing a raise in downstream products *S*-sulfocysteine (SSC), mediating excitotoxicity, and thiosulfate, a catabolic intermediate/product of H_2_S metabolism. Here, we report a full-body knock-out mouse model for SOX deficiency (SOXD) with a severely impaired phenotype. Amongst the urinary biomarkers, thiosulfate showed a 45-fold accumulation in SOXD mice representing the major excreted S-metabolite. Consistently, we found increased plasma H_2_S, which was derived from sulfite-induced release from persulfides as demonstrated *in vitro* and *in vivo*. Mass spectrometric analysis of total protein persulfidome identified a major loss of persulfidation in 20% of the proteome affecting enzymes in amino acids and fatty acid metabolism. Urinary amino acid profiles indicate metabolic rewiring suggesting partial reversal of the TCA cycle thus identifying a novel contribution of H_2_S metabolism and persulfidation in SOXD.

## Introduction

Sulfite oxidase (SOX) deficiencies are life-threatening disorders of sulfur metabolism, impacting the terminal step of cysteine degradation: the oxidation of toxic sulfite (SO_3_^2-^) to sulfate (SO_4_^2-^, Figure 1A). SOX is a molybdenum- and heme containing dimeric enzyme localized to the intermembrane space of mitochondria where it transfers electrons derived from sulfite oxidation to cytochrome c thus contributing to mitochondrial respiration. SOX deficiency may either be caused by pathogenic variants in the *SUOX* gene encoding for the SOX protein (Claerhout et al., 2018; Tan et al., 2005) resulting in the so-called isolated SOX deficiency (SOXD) or by defects in the biosynthesis of its molybdenum cofactor (Moco), which is synthesized by a multistep pathway (Mayr et al., 2021) leading to Moco deficiency (MoCD). Both disorders, SOXD and MoCD, are symptomatically similar, underlying the importance of SOX. Patients with SOXD and MoCD display neonatal feeding difficulties, pharmaco-resistant seizures, severe white matter abnormalities, enlarged ventricles, subcortical cavities, microcephaly, and other dysmorphic changes (Carmi-Nawi et al., 2011; Claerhout et al., 2018; Tan et al., 2005; van der Knaap & Valk, 2001). Patients with MoCD exhibit in addition also xanthine urolithiasis due to secondary xanthine oxidase deficiency.

**Figure 1:**
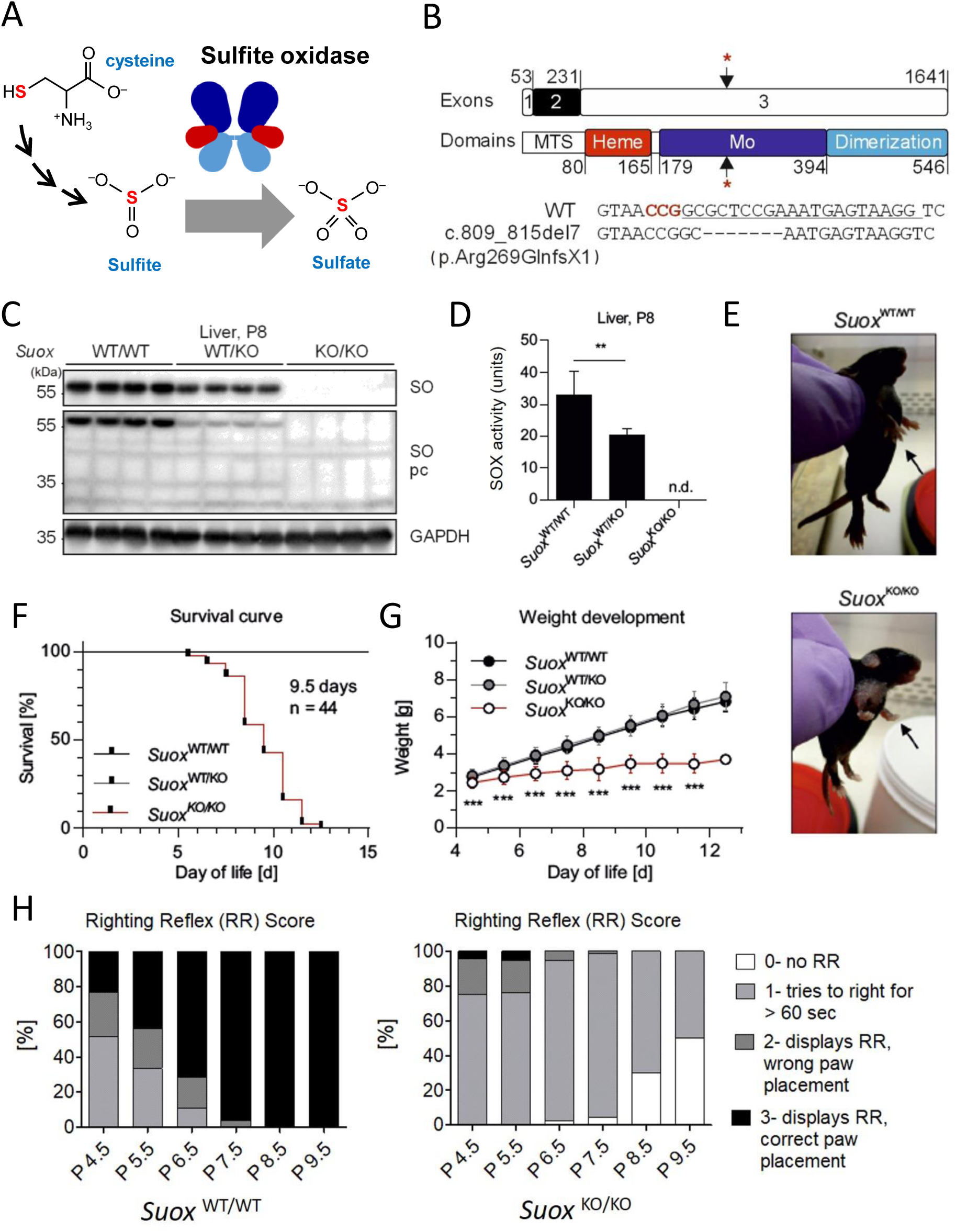
Generation and characterization of *Suox*-knockout in C57BL6/J mice. **(A)** Schematic representation of the catalytic function of sulfite oxidase. **(B)** Schematic representation of murine *Suox* exon structure and SOX domain structure. Asterisks indicate CRISPR/Cas9 targeting site. **(C)** Western blot for SOX in liver lysates of littermates from *Suox*^WT/KO^ x *Suox*^WT/KO^ breedings using two different antibodies (*n* = 4). **(D)** Cytochrome *c*-SOX activity detected in liver lysates of littermates from *Suox*^WT/KO^ x *Suox*^WT/KO^ breedings (*n* = 8). Student’s t test was performed as indicated. p value: ** < 0.01; * < 0.05; ns > 0.05. **(E)** Phenotype of *Suox*^WT/WT^ and *Suox*^KO/KO^ mice at P9. Display of potential front paw spasticity and stunted growth in *Suox*^KO/KO^ mice (bottom) compared to *Suox*^WT/WT^ mice (top). **(F)** Kaplan-Meier plot for survival of *Suox*^KO/KO^ mice (n = 28) compared to *Suox*^WT/WT^ (n = 25) and *Suox*^WT/KO^ (n = 60) littermates. **(G)** Weight development of all genotypes of the SOX-deficient line, starting at P4.5. Asterisks indicate difference of *Suox*^KO/KO^ mice to Suox^WT/WT^ and *Suox*^WT/KO^mice. **(H)** Overtime righting reflex evaluation according to criteria modified from DiDonato and Bogdanik for *Suox*^WT/WT^ and *Suox*^KO/KO^ mice. Testing was performed at least three times per animal and day (technical replicates), with at least four animals per genotype and day (biological replicates).

Sulfite was early recognized to be accumulated in urine of patients (Mudd et al., 1967) and later shown to be a reactive, cell-toxic agent (Zhang et al., 2004). We have shown that the accumulating sulfite-cystine adduct, *S*-sulfocysteine (SSC), is key causative factor of neurodegeneration in the disease (Kumar et al., 2017). SSC was found to act as an NMDA receptor agonist, whose effect could be blocked by the application of the NMDA receptor antagonist memantine, inhibiting phenotype development in a pharmacologically induced mouse model of MoCD (Kumar et al., 2017). In addition, SSC (together with S-sulfohomocysteine) may also inhibit gamma-glutamylcysteine synthetase (Moore et al., 1987). As a result of SSC accumulation, cysteine/cystine and homocysteine were found to be depleted in SOXD/MoCD patients suggesting major changes in the overall cysteine homeostasis.

A third accumulating biomarker in SOXD/MoCD is thiosulfate, a catabolic end product of sulfur metabolism which is derived from H_2_S. H_2_S is synthesized by various enzymes following non-canonical reactions in the reverse transsulfuration pathway (cystathionine β-synthase, CBS and cystathionine γ-lyase, CSE). In addition, mercaptopyruvate sulfur transferase forms a protein-bound persulfide also contributing to H_2_S formation (Kohl et al., 2019). H_2_S is an important signaling molecule with versatile biological implications. The two most well studied mechanisms of its actions are i) interactions with metalloproteins (Domán et al., 2023) and ii) redox reactions with thiol proteins (Filipovic et al., 2018; Nagy et al., 2014). H_2_S, cysteine and thiosulfate metabolic pathways are tightly linked to cysteine persulfidation. This posttranslational modification was reported to have pivotal role by regulating and protecting functions on thiol proteins (Dóka et al., 2020; Mustafa et al., 2009; Zivanovic et al., 2019).

Interestingly, thiosulfate also accumulates in ethyl malonic encephalopathy (EE) caused by defects in the enzyme persulfide dioxygenase encoded by the *ETHE*1 gene. This defect results in mitochondrial dysfunction, neurodegeneration and childhood death (Tiranti et al., 2004). Targeted sulfur metabolome profiling in humans suffering from ultrarare enzyme deficiencies of sulfur metabolic pathways was reported recently, which highlighted that the homeostasis of sulfur metabolites largely rely on catabolic pathways while H_2_S-synthesizing routes can efficiently compensate for each other’s activity (Kožich et al., 2022).

To date, there are three mouse models recapitulating the different forms of MoCD (type A, B, and C) generated by whole body gene knock outs in *Mocs1*, *Mocs2* and *gphn* genes (Feng et al., 1998; Jakubiczka-Smorag et al., 2016; Lee et al., 2002), all presenting with a severe phenotype causing death at the age 1-8 days. In addition, mice treated with high dosages of tungstate represent a pharmacological model for MoCD (Kumar et al., 2017). All models have in common, that in addition to the lack of SOX activity, also other Moco-containing enzymes are dysfunctional and in case of xanthine oxidoreductase deficiency, accumulating xanthine deposits as kidney stones that may trigger acute kidney failure (Jakubiczka-Smorag et al., 2016) potentially contributing to the lethality in MoCD mice.

Here we have generated an animal model for SOXD using the CRISPR/Cas9 genome editing to disrupt the murine *Suox* gene. We report the phenotypic characterization and detailed biomarker profile that enabled us to identify a novel contribution of H_2_S biology in the underlying pathomechanism of SOX deficiency. We found a sulfite-dependent liberation of H_2_S from persufides leading to massive metabolic alterations in SOX-deficient mice.

## Results

### *Suox*^KO/KO^ mice show a severe phenotype without sign of neurodegeneration

Using a CRISPR/Cas9 approach we generated a mouse line with a 7-bp deletion in the *Suox* gene starting at position c.809 triggering a frameshift, resulting in a stop codon at position p.Arg269GlnX1 of SOX protein (Fig. 1B). The resulting off-springs segregated as expected in a 1:2:1 manner into *Suox*^WT/WT^, *Suox*^WT/KO^, *Suox*^KO/KO^ suggesting no lethality during embryo development (Suppl. Fig. 1). In liver extracts, the organ with highest SOX expression (Belaidi et al., 2015) we found neither SOX protein nor any detectable SOX activity in *Suox*^KO/KO^, while in *Suox*^WT/KO^ approx. 50% reduced SOX expression was observed (Fig. 1C-D), confirming the successful deletion of the enzyme in knockout animals *Suox*^KO/KO^.

Phenotypically, *Suox*^KO/KO^ animals displayed delayed fur and whisker development (Fig. 1E), similar to *Mocs1*- and *Mocs2*-deficient mice (Jakubiczka-Smorag et al., 2016). Survival of *Suox*^KO/KO^ mice was significantly impaired, with a lifespan ranging from 5 to 12 days and a median survival of 9.5 days (Fig. 1F), which was longer by 1-2 days when compared to *Mocs1*- and *Mocs2*-deficient animals (Jakubiczka-Smorag et al., 2016; Lee et al., 2002) suggesting a minor contribution of the other Moco-dependent enzymes to the pathology of MoCD mice (Jakubiczka-Smorag et al., 2016; Lee et al., 2002). *Suox*^KO/KO^ animals showed postnatal milk spots, but failed to thrive starting from P4. Weight gain was significantly impaired following day 4, and was halted or started to decline one day before death or reaching a score requiring killing (Fig. 1G). Mice used for subsequent analysis were sacrificed at day 8, if not stated otherwise.

Given the neurodegenerative phenotype of human patients (Carmi-Nawi et al., 2011; Claerhout et al., 2018; Tan et al., 2005; van der Knaap & Valk, 2001), we first analyzed mice heads by magnetic resonance imaging. *Suox*^KO/KO^ brains appeared round, while the shape of their WT littermates was elliptical and total brain volume of *Suox*^KO/KO^ mice was decreased by 29.5% when Compared to *Suox*^WT/WT^ (Suppl. Fig. 2A-B). Immunohistochemistry for microtubule-associated protein 2 (MAP-2) as well as for the neuronal nuclear antigen, NeuN, revealed no differences between *Suox*^KO/KO^ mice and their WT littermates as judged by a quantitative analysis in motor cortex layer V (Suppl. Fig. 2C-D). However, when we assessed motor skill development using surface righting reflex testing at P4.5, *Suox*^KO/KO^ animals did not score strikingly different from *Suox*^WT/WT^ littermates while motor skill development appeared halted on P5.5 and worsened on subsequent days of testing. At P9.5, approximately 50 % of surviving *Suox*^KO/KO^ animals lost any righting reflex, while the remaining animals attempted for longer than 60 sec (Fig. 1H). In comparison, by P8.5 all WT littermates performed the reflex with correct paw placement (Fig. 1H). In conclusion, we found that *Suox*^KO/KO^ present with a severe phenotype, short live span and arrest in motor skill development while no signs of morphological changes in the brain were observed within the first eight days of life.

### Thiosulfate is the major S-metabolite excreted in *Suox*^KO/KO^ mice

MoCD and SOXD patients display increased urinary sulfite, S-sulfocysteine (SSC), S-sulfohomcysteine, and thiosulfate (Kožich et al., 2022) (Fig. 2A). To understand the flux and organ contribution of these biomarkers, we determined their levels in brain, liver and kidney as well as in plasma and urine of *Suox*^KO/KO^ mice and control mice (Fig. 2B-D, Fig. 2G-I, and Suppl. Fig. 3) and compared these with total cysteine (Fig. 2E) and bioavailable H_2_S levels in plasma (Fig. 2F).

**Figure 2:**
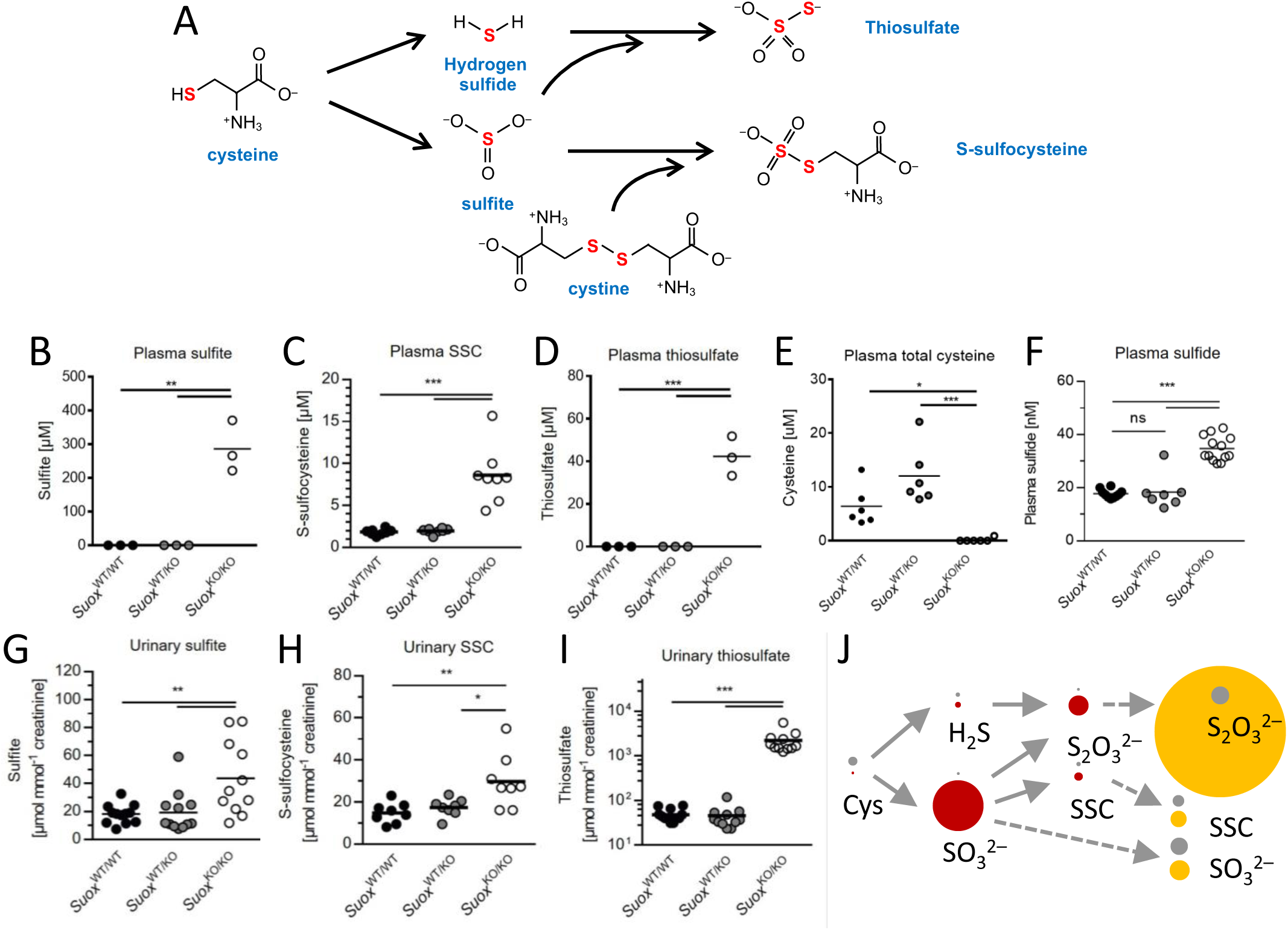
Levels of main biomarkers of SOXD and metabolites of cysteine catabolism in *Suox* mice. **(A)** Schematic representation of cysteine catabolism. Levels of highlighted metabolites (Sulfite, S-sulfocysteine, thiosulfate, and hydrogen sulfide) and free cysteine have been determined in this study. **(B)** Determination of plasma sulfite levels in *Suox* mice. **(C)** Determination of plasma S-sulfocysteine levels in *Suox* mice. **(D)** Determination of plasma thiosulfate levels in *Suox* mice. **(E)** Determination of plasma free cysteine levels in *Suox* strain mice. **(F)** Determination of plasma hydrogen sulfide levels in *Suox* mice. **(G)** Determination of urinary sulfite levels in *Suox* mice. **(H)** Determination of urinary S-sulfocysteine levels in *Suox* mice. (I) Determination of urinary thiosulfate levels in *Suox* mice. One-way ANOVA with Tukey’s post-hoc test for pairwise comparisons was performed as indicated. *p* value: *** < 0.001; ** < 0.01; * < 0.05; ns > 0.05. **(J)** Cartoon highlighting the flux of sulfur-containing metabolites in *Suox*^KO/KO^ mice (colored circles; red circles: plasma; yellow circles: urine) in comparison to WT mice (gray circles). Circles are drawn in scale to their respective concentrations in plasma and urine, with the exception for H_2_S, which is 10-fold enlarged.

Sulfite was found to accumulate in all three organs, with liver showing the highest sulfite level as expected for the major SOX-expressing organ (Suppl. Fig. 3A). In plasma, sulfite showed highest level of accumulation reaching nearly 300 µM (Fig. 2B), while urinary sulfite (Fig. 2G) was only moderately increased (2-fold) suggesting a poor clearance. For SSC, we found no alteration in liver and kidney, a minor increase in brain (1.6-fold), while in plasma, SSC levels were strongly elevated (4-fold of WT) in plasma reaching 8.6±3.4 µM in plasma. In contrast, urinary excretion was again only 2-fold increased, when compared to WT, which is much lower than levels in human patients that show strong accumulation in urine ranging from 100 to 500 µmol SSC/mmol creatinine (Kožich et al., 2022). Levels of SSC in plasma mirrored the drop in total plasma free cysteine in *Suox*^KO/KO^ mice, which was not detectable while WT mice showed 6.3 ± 2.2 µM free cysteine. Based on these data we conclude, that the SSC level reached saturation due to a quantitative conversion of cystine by accumulating sulfite (Jakubiczka-Smorag et al., 2016; Kožich et al., 2022).

For thiosulfate, as a biomarker of H_2_S catabolism, we found an accumulation in all three organs with highest levels detected in kidney (Suppl. Fig. 3C). Total thiosulfate in plasma reached 42.3 ± 4.4 µM being 5-times higher than SSC but much lower than circulating sulfite. To our great surprise, thiosulfate excretion was the highest of all biomarkers representing 45-fold more than observed in WT mice. The total excretion of 2,200 µmol thiosulfate/mmol creatinine exceeds that of sulfite by almost 40-fold suggesting that the vast majority of excreted sulfur is derived from H_2_S and related species. Finally, we measured bioavailable H_2_S in plasma (Ditrói et al., 2019) and found a significant two-fold increase in *Suox*^KO/KO^ mice clearly indicating an alteration in H_2_S metabolic pathways (Fig. 2F). Taken together, all biomarker analyses collectively identified a major route for sulfur excretion from accumulating sulfite into thiosulfate (Fig. 2J).

Given the metabolic link between thiosulfate and H_2_S, we next investigated H_2_S biogenesis, homeostasis and catabolism in *Suox*^KO/KO^ mice. In a proteome-wide analysis (Suppl. Fig. 4) we inspected levels of enzymes involved in H_2_S biogenesis and found no biologically relevant alteration for cystathionine β-synthase (CBS), cystathionine γ-lyase (CSE), and 3-mercaptopyruvate sulfurtransferase in liver and kidney thus excluding increased H_2_S biosynthesis as sources of H_2_S and thiosulfate (Suppl. Fig. 4). Sulfide:quinone oxidoreductase (SQR), persulfide dioxygenase (PDO), and thiosulfate sulfurtransferase (TST) catalyze individual steps in the H_2_S oxidative pathway producing glutathione and other low-molecular weight persulfides that are further converted into sulfite and thiosulfate (Hildebrandt et al., 2013; Libiad et al., 2014). Again, no biologically relevant change was observed for those proteins. Out of all the proteins involved in cysteine metabolism there was only one enzyme showing a significant change; cysteine dioxygenase (Cdo1), the first enzyme in oxidative cysteine catabolism, was 7.3-fold reduced being in-line with the loss of free cysteine in plasma (Dominy et al., 2006). Therefore, our observed increase in H_2_S and the massive excretion of thiosulfate should have a different origin, which we investigated next.

### Sulfite induces a release of H_2_S from persulfides

In light of the accumulation of sulfite observed in all investigated organs with highest concentration in plasma, we raised the question whether sulfite reacts with persulfidated small molecules and proteins known to contribute significantly to the overall pool of labile H_2_S in the organism (Ono et al., 2014; Zivanovic et al., 2019). We hypothesized that sulfite treatment of persulfides could result in the formation of S-sulfonylated species with the concomitant release of H_2_S (Fig. 3A) (Zivanovic et al., 2019). First, we tested the reactivity of sulfite towards persulfides using an H_2_S electrode and various low-molecular weight and protein persulfide models. *N*-acetylpenicilinamine persulfide (NAP-SSH) was reported to display slow H_2_S dissociation kinetics at pH 7.4 (Artaud & Galardon, 2014). When adding of 20-fold molar excess of sulfite (1 mM) to 50 µM NAP-SSH, the dissociation rate of H_2_S was greatly increased within seconds leading the rapid re-equilibration (Fig. 3B). Next, we investigated two different protein model substrates previously reported to be persulfidated at key cysteine residues (Cuevasanta et al., 2015), human serum albumin (HSA, Fig. 3C) and glycerol-3-phosphate dehydrogenase (GAPDH, Fig. 3D) (Dóka et al., 2016; Wedmann et al., 2016; Zhang et al., 2014) . Similar to NAP-SSH, again sulfite-dependent H_2_S release was observed with a rapid H_2_S spike and re-equilibration with GAPDH (Fig. 3D) collectively confirming a non-enzymatic release of H_2_S from various persulfidated targets.

**Figure 3.**
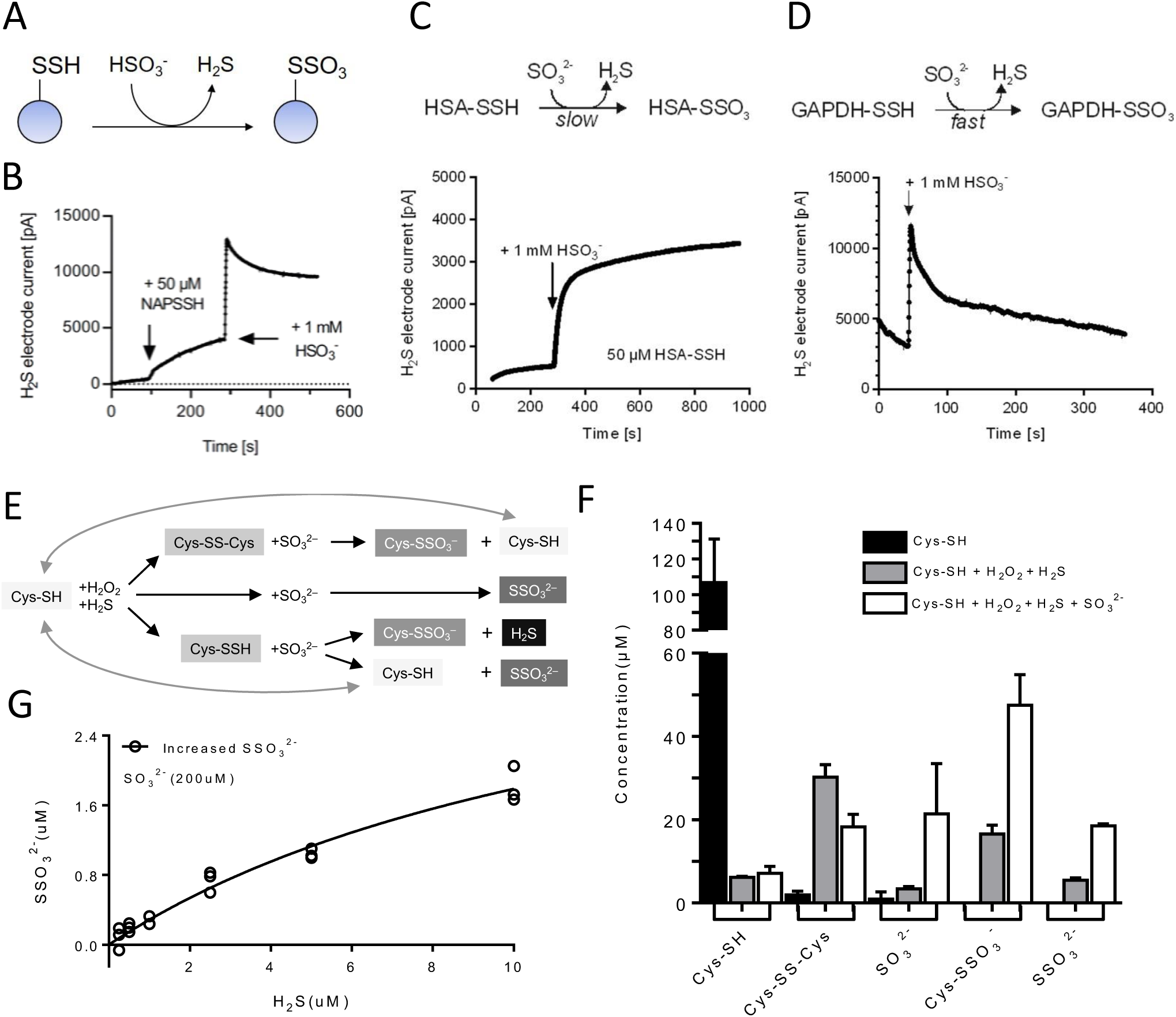
Sulfite triggers protein depersulfidation and releases hydrogen sulfide. **(A)** Schematic reaction of persulfides with HSO_3_^-^ at pH 7.4, producing persulfonates (*S*-sulfonates) and releasing H_2_S. **(B)** Representative reaction of 50 µM NAP-SSH with 1 mM 28 HSO ^-^ at pH 7.4, measuring H S release. **(C)** H S electrode signal of 50 μM HSA-SSH in 50 mM PBS, pH 7.4, reacting with a 20-fold molar excess of sulfite (n = 3). **(D)** H_2_S electrode signal of 50 μM GAPDH-SSH in 50 mM PBS, pH 7.4, reacting with 20-fold molar excess of sulfite (n = 3). **(E)** Schematic representation of cysteine persulfides depersulfidated by sulfite and formed H_2_S. **(F)** End-product measurements of *in vitro* interaction between sulfite and cysteine persulfides. Error-bars indicate standard deviation. **(G)** H_2_S-depending of non-enzymatic thiosulfate formation with high-level sulfite.

The reaction of sulfite with persulfides also displays a path towards S-sulfonylation other than the oxidation of persulfides (Dóka et al., 2020; Zivanovic et al., 2019). In order to monitor this half of the reaction and to answer the question to which extent sulfite reacts with the outer sphere sulfur of the persulfide leading to thiosulfate and free thiol release, we designed another *in vitro* experiment (Fig. 3E). As described before, mild oxidation of cysteine to sulfenic acids increased the reactivity of thiols towards H_2_S thus forming cysteine persulfides *in vitro* (Zivanovic et al., 2019).

We treated 100 µM cysteine with equimolar amounts of H_2_O_2_ and H_2_S for 1 hour and determined the amount of cysteine, cystine, sulfite, SSC, and thiosulfate (Fig. 3F). As expected, we found 30 µM cystine, representing 60% of cysteine being fully oxidized by H_2_O_2_. However, approx. 18 µM SSC and 7 µM thiosulfate were also formed, leaving only ∼15% of the remaining cysteine to be potentially persulfidated. When treating this mixture with 100 µM sulfite, ∼ 10 µM of cystine was depleted resulting in the formation of SSC and cysteine. The fact that 30 µM additional SSC was formed, suggests that cysteine-persulfide donated a significant fraction (approx. 10 µM) to this reaction. Given that only 20 µM remaining sulfite was detected, suggests that a large fraction of sulfite was bound in SSC. In addition, we found an increase in thiosulfate formation that could either suggest an abstraction of the persulfide’s outer sulfur by sulfite and/or an H_2_S-dependent formation of thiosulfate with sulfite. We have shown the existence of the latter reaction by titrating sulfite with substoichiometric amounts of H_2_S leading to a thiosulfate-to-H_2_S ratio of approx. 1:4 in the presence of sulfite excess (200 µM, Fig. 3G), conditions representing levels in *Suox*^KO/KO^ mice. In aggregate, our *in vitro* studies demonstrate that at pathological sulfite concentrations, various persulfides release H_2_S resulting in the formation of S-sulfonylated reaction products.

### Sulfite changes the persulfidome of *Suox*^KO/KO^ mice

Following our findings that sulfite reacts with persulfides, we wanted to test how accumulation of sulfite affects the global persulfidome landscape. Using the dimedone switch method (Zivanovic et al., 2019) for selective persulfide labeling (Fig. 4A) we first observed, in-gel, that liver samples from *Suox*^KO/KO^ mice show significantly lower protein persulfide levels (Fig. 4B-C). To quantify this change and understand its biological effect we next performed a label-free persulfidome analysis and identified 2,457 persulfidated proteins in both WT and *Suox*^KO/KO^ mouse liver samples (Table S1). Amongst those, 571 proteins passed the significance threshold (presence in 3 out 4 samples, at least 30% change and p> 0.05) of which 441 proteins showed clear decrease in protein persulfidation (Fig. 4D-E, Suppl. Fig. 5A-B). To exclude that this change originates from the change in total protein level, we also performed label-free total proteome analysis (Suppl. Fig. 5C-D) and double plotted the persulfidome change to the total proteome change (Suppl. Fig. 5E). Indeed, majority of identified proteins seems to be unaffected by the change in protein expression levels.

**Figure 4:**
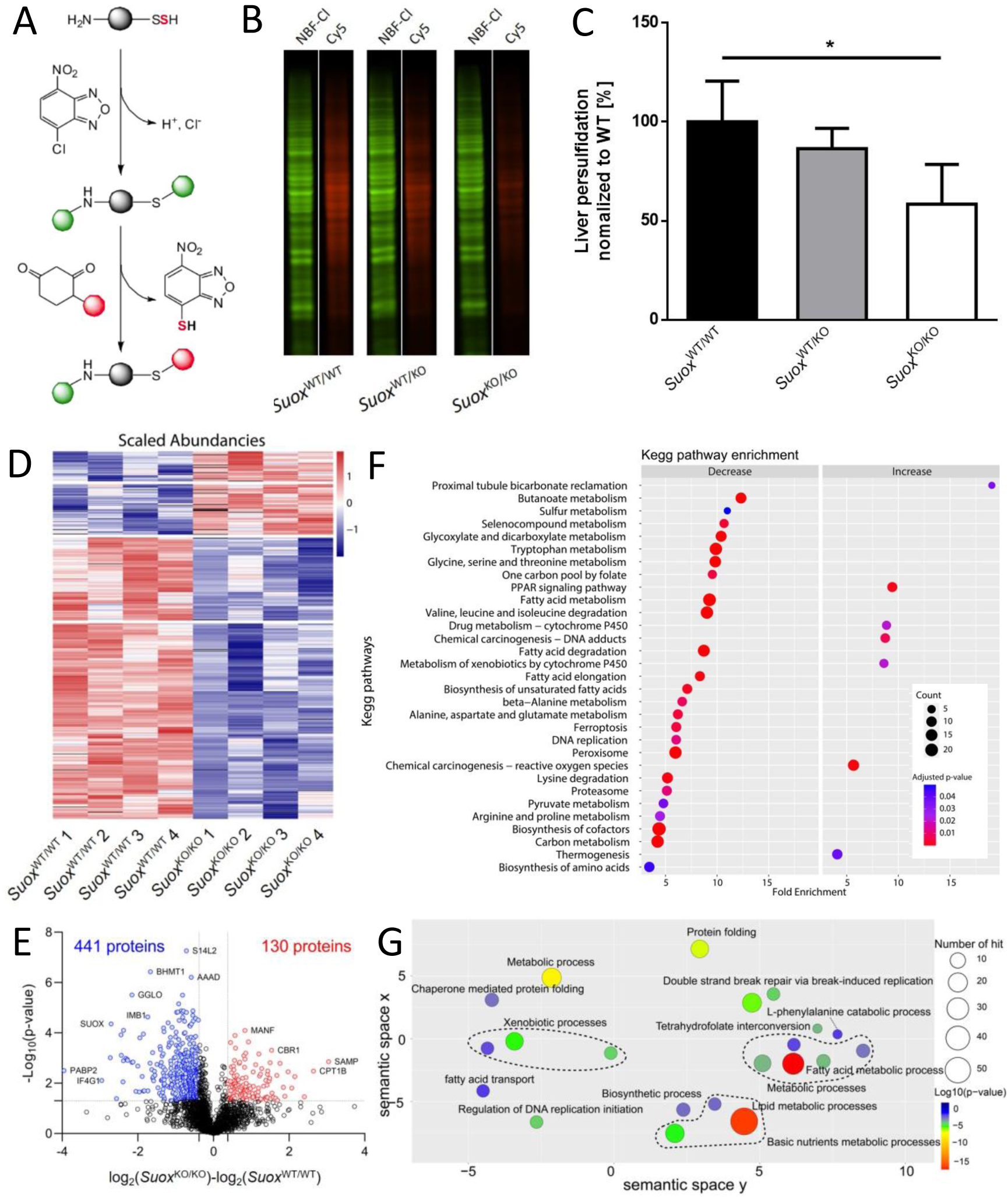
Persulfidome remodeling in *Suox*^KO/KO^ mice. **(A)** Setup for the detection of protein-bound persulfides in the assay described by Zivanovic et al. Crude extract from liver is treated with NBF-Cl, which reacts with persulfides, free thiols, sulfenic acids and amine groups. The NBF-blocked persulfides are nucleophillically attacked by the DAz-2 moiety, which in turn is cross-linked to a fluorescent Cy5 moiety via click-chemistry. **(B)** Representative Cy5 (persulfonates signal) and NBF-Cl (control signal) fluorescence signals measured by a fluorescence scanner. (Cy5 –fire scale; NBF-Cl –green). **(C)** Quantification of a specific rectangular area identical for all lanes. All Cy5 signals were normalized to their respective NBF-Cl signals, and further normalized to the mean of *Suox*^WT/WT^ signals (n = 3). Error-bars indicate standard deviation. One-way ANOVA was performed as indicated. *p* value: *** < 0.001; ** < 0.01; * <0.05; ns > 0.05. **(D)** Heatmap showing the significant changes of protein persulfidation in *Suox*^KO/KO^ mouse livers compared to wild type animals (Welch’s test, p<0.05). **(E)** Volcano plot depicting statistical significance plotted against the log_2_-fold change of persulfidated proteins in *Suox*^KO/KO^ mouse liver relative to wild type animals. Significance was established using Welch’s t-test (two-sided), with a p-value threshold of < 0.05. Fold change cut-offs were established at 30%. The numbers of proteins with significantly altered persulfidation states are indicated, **(F)** Kegg pathway enrichment analysis using DAVID. The graph shows the top 30 significant (Benjamini adjusted p-value < 0.01) and most enriched terms, with color gradient signifying the adjusted p-value and circle size the number of proteins. **(G)** GO (Biological process) term enrichment analysis of the 441 proteins found to have decreased persulfidation levels in *Suox*^KO/KO^ mice. REVIGO was used to plot the enrichment analysis performed in DAVID. Circle dimensions denote the protein count within specific GO terms, while color gradients depict the degree of significance. Similar biological processes are grouped together with dashed-line.

Kegg pathway enrichment analysis of proteins that showed significant decrease in protein persulfidation suggests that a variety of metabolic processes is affected, such as amino acid metabolic pathways, lipid metabolism, pyruvate metabolism, but also ferroptosis, DNA replication, PPAR signaling etc (Fig. 4F). A complementary GO term enrichment analysis identified similar pathways (Fig. 4G).

Based on our *in vitro* studies with persulfidated cysteine, a decrease in protein persulfidation should be mirrored by an increase in S-sulfonylation. However, no method for selective enrichment of this modification exists today. Nonetheless, considering that the first step of the dimedone-switch method is blocking all thiols, persulfides, sulfenic acids and amines, preventing any artificial persulfide oxidation, we looked for the S-sulfonylated peptides in those samples. We could detect 45 peptides passing the inclusion criteria (being present in at least 4 out of 5 samples) corresponding to 40 proteins, of which 40 % showed significant increase in *Suox*^KO/KO^ mice (peptides normalized to actual protein levels, 1.3-fold change) (Suppl. Fig. 6). Despite the low number of identified targets, both Kegg pathway and GO term enrichment analyses pointed towards metabolic processes, such as lipid and amino acid metabolism, as well as ferroptosis, being very similar to the results of the corresponding enrichments analyses in our persulfidation data set.

### Changes in the amino acid profile suggest metabolic rewiring

We have conclusively demonstrated that sulfite toxicity in mice resulted in H_2_S release from S-persulfidated stores leading to a massive excretion of thiosulfate. H_2_S toxicity has been attributed to the inhibition of mitochondrial respiration due to the blockage in complex IV (Tiranti et al., 2009). In a recent study by Banerjee and coworkers it was shown that H_2_S excess may cause reversal of the respiratory chain, an increase in mitochondrial NADH pool thus leading to a partial reversal of the TCA cycle with increased glycolysis to generate the required ATP (Carballal et al., 2021). In order to feed the TCA cycle, increased glutaminolysis has been demonstrated as major source for α-ketoglutarate (Carballal et al., 2021).

To get further insights into the metabolic changes in *Suox*^KO/KO^ mice, we determined the amino acid profile in urine (Fig. 5). Remarkably, levels of nearly all amino acids were decreased in *Suox*^KO/KO^ mice, with cystine and glutamine showing the strongest reduction (Fig. 5). While cystine depletion is directly related to quantitative conversion to SSC, an over 10-fold reduction in glutamine supports the above-mentioned consequences of H_2_S toxicity on mitochondrial respiration. In addition, all other catabolic amino acid precursors (Glu, His, Arg, Pro) of α-ketoglutarate were also reduced (Fig. 5). Furthermore, amino acids contributing to acetyl-CoA and pyruvate formation (Ala, Ser, Gly, Thr, Cys) were also depleted, collectively arguing for a metabolic arrest in *Suox*^KO/KO^ mice. This is in line with the persulfidome data and the observation that animals stop gaining weight at day 4 suggesting a catabolic state of metabolism. Taken together, SOX-deficient mice disclosed an unexpected rewiring of cysteine catabolism into the H_2_S pathway caused by a sulfite-induce de-persulfidation, H_2_S release providing a new view on this ultra-rare and devastating metabolic disorder.

**Figure 5:**
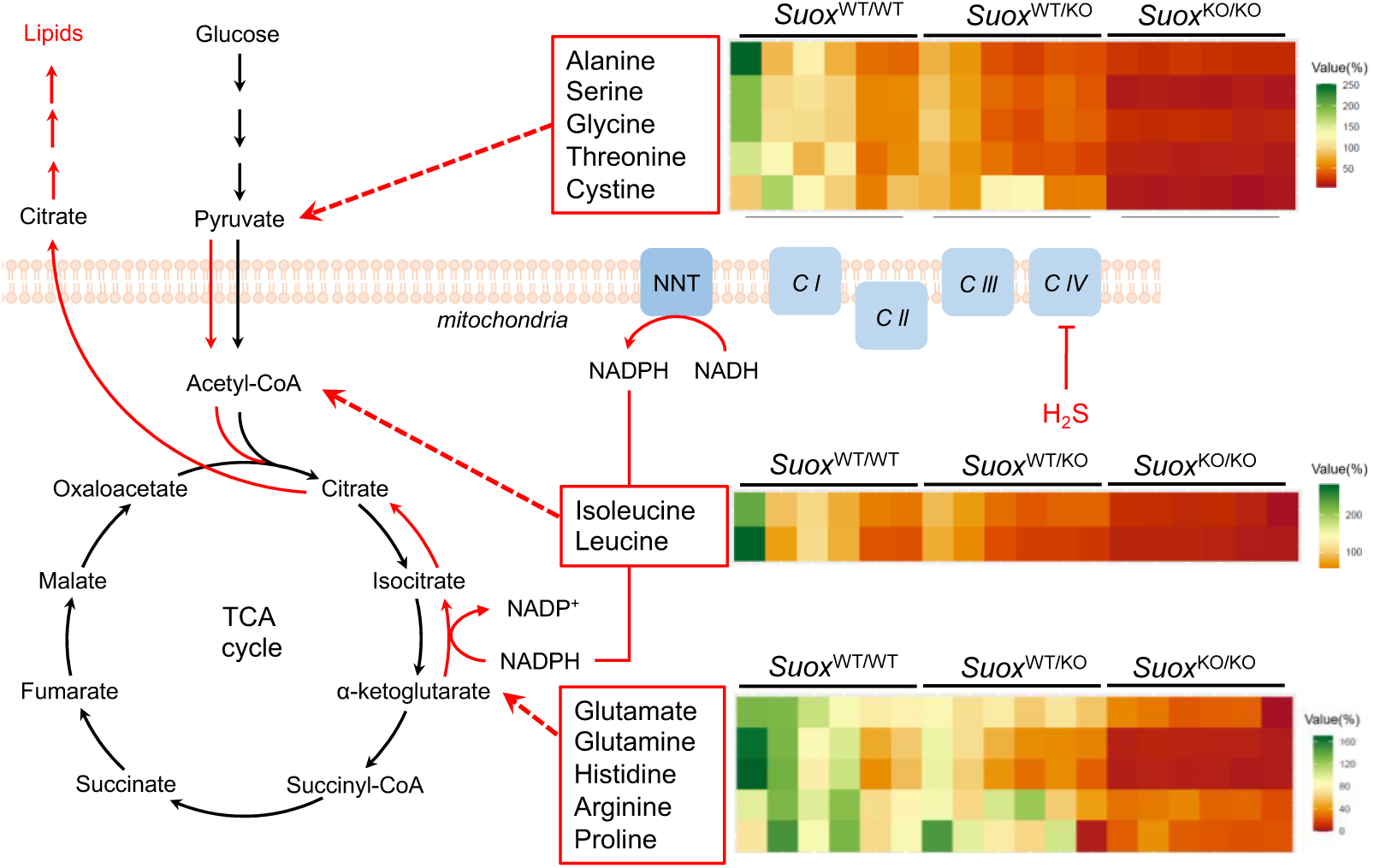
Metabolic impacts of elevated H_2_S in *Suox*^KO/KO^ mice. Schematic representation of partial reversal of TCA cycle and relative changes (n=6) of urinary amino acids in comparison to ***Suox*^WT/WT^**. Amino acids are group according to their contribution as catabolic precursors of α-ketoglutarate, acetyl-CoA and pyruvate.

## Discussion

Based on the symptomatic similarity between MoCD and SOXD, defective SOX enzyme is recognized as the major cause of death in both diseases. So far, only MoCD mice models were described, thus, we generated and characterized the first SOXD mouse model in this study. Human patients with SOXD were reported with feeding difficulties, unprovoked and drug-resistant seizures shortly after birth, and further developed secondary pathologies such as lens dislocation, neurodegeneration with cysts formation, and microcephaly leading to a mean survival of four years. Remarkably, MoCD/SOXD patients are often misdiagnosed with hypoxic ischemia encephalopathy. In comparison, *Suox*^KO/KO^ mice were observed with severe growth retardation as they stopped gaining weight from postnatal day 4, which resulted in a mean lifespan of 9.6 days.

Despite that *Suox*^KO/KO^ mice die in young age, no neurodegeneration was found in those mice. In addition, in previously reported MoCD mouse models – neither in *Mocs*1-(Lee et al., 2002) nor *Mocs*2-deficient (Jakubiczka-Smorag et al., 2016) mice – any neurodegeneration had been reported. This observation may have the following reasons: (i) Brain development in newborn rodents is not as complete as in human infants. Rodent brains at postnatal days 7-10 are equivalent to human infant brains (Semple et al., 2013). Given that *Suox*^KO/KO^ mice die within 9.6 days; we expect to be outside the developmental window of the comparable age of human patients. (ii) Urinary SSC levels in mice were at 30 µmol/mmol creatinine thus being much lower than in human patients (129 µmol/mmol creatinine) (Kožich et al., 2022), suggesting a reduced overall production and clearance rate of SSC. Given that plasma cysteine was depleted in *Suox*^KO/KO^ mice, we conclude that limiting cystine concentrations can be considered the primary cause of lower SSC levels in mice. Therefore, we considered other alterations in sulfur-containing metabolites as drivers for the disease pathology in mice.

When comparing the excretion between sulfite (45 mmol/mol creatinine), SSC (32 mmol/mol creatinine), and thiosulfate (2,200 mmol/mol creatinine) it becomes clear that the vast majority (97%) of sulfur is excreted as thiosulfate, the catabolic oxidation product of H_2_S and sulfite. Thus, we wondered if H_2_S and thiosulfate also contributed to the SOXD pathomechanism. Patients with two other inherited metabolic disorders, sulfide:quinone oxidoreductase deficiency (SQORD) and ethylmalonic encephalopathy (EE), were also reported with approximately two-fold increased levels of H_2_S, but only EE patients showed thiosulfate elevation comparable with MoCD/SOXD patients in plasma (Kožich et al., 2022). SQORD is caused by the mutations in the *SQOR* gene encoding for a mitochondrial enzyme that oxidizes H_2_S to persulfides (Jackson et al., 2012; Lagoutte et al., 2010). EE patients, on the other hand, harbor mutation in the *ETHE1* gene, which encodes for PDO enzyme converting persulfides into thiols and sulfite (Kabil & Banerjee, 2012). Under conditions of PDO deficiency, accumulating persulfides are further metabolized in a sulfite-dependent manner by thiosulfate sulfurtransferase (TST) yielding thiosulfate. Remarkably, similar to MoCD and SOXD, patients with SQORD or EE suffer from brain lesions, which are considered the consequence of H_2_S accumulation.

H_2_S is gaining growing interest as gasotransmitter due to its role in mediating persulfidadtion of proteins and small molecules with beneficial effects on stress resistance and longevity (Zivanovic et al., 2019). However, at high concentrations, H_2_S binds to cytochrome *c* oxidase thus suppressing mitochondrial oxidative phosphorylation as it was shown in fibroblasts derived from SQORD and EE patients (Friederich et al., 2020; Grings et al., 2019). In contrast to our SOXD mouse model, SQORD and EE mice (Marutani et al., 2021; Tiranti et al., 2009) show milder disease progressions with average lifespans of weeks to months at comparable levels of H_2_S. *Sqor*^-/-^ mice were indistinguishable from wild-type littermates before weaning, but later developed ataxia resulting in premature death in an age of 10 weeks. On the other hand, *Ethe1*^-/-^ mice have a more severe phenotype than *Sqor*^-/-^ mice, given that they showed growth arrest from P15 and died between five to six weeks of age. In addition, a large and comparative study that included several sulfur-metabolic enzyme disorders in humans showed that in addition to MoCD/SOXD patients, also EE showed moderate but significant accumulation of sulfite (Kožich et al., 2022). In aggregate, our findings suggest that SOXD resulted in a massive accumulation of both sulfite and thiosulfate in mice leading to a collective pathology of sulfite and H_2_S metabolites, given that thiosulfate has been shown to be rather safe (Kumar et al., 2017) and even protective (Macabrey et al., 2022; Zhang et al., 2021).

Our observed significant increase in plasma H_2_S in *Suox*^KO/KO^ mice is comparable with that seen in EE patients (Kožich et al., 2022), however, none of the enzymes involved in H_2_S biosynthesis or catabolism were found to be altered. Therefore, we hypothesized that there were other sources of H_2_S-derived thiosulfate. Given that sulfite attacks various forms of oxidized cysteines, as reported for cystine-dependent SSC formation (Kumar et al., 2017), we demonstrated here that sulfite also cleaves the S-S bond of persulfidated cysteines, either in the metabolite cysteine-persulfide or in peptides and proteins. The chemical properties of persulfides are unique in the sense that the inner and the outer sulfurs are not equivalent in their reactivity. In comparison, the pK_a_ values of RSH are approx. 8-10 while the pK_a_ value of H_2_S is approx. 7, which makes HS^-^ a better-leaving group than RS^-^ (Saund et al., 2015). To study which sulfur atom is the main site for the nucleophilic attack of sulfite and which reaction products are formed, we treated persulfidated NAP, HSA, and GAPDH with sulfite and observed different kinetics for H_2_S release. When using *in vitro*-generated Cys-SSH, sulfite treatment resulted in a near stoichiometric formation of S-sulfocysteine. In addition, a minor formation of thiosulfate was observed due to H_2_S- and H_2_O_2_-dependent reaction with sulfite. Therefore, we conclude that sulfite-induced release of H_2_S from persulfidated stores leads to an acute H_2_S intoxication and loss of protein persulfidation. Consistently, our proteome-wide study in liver extracts of *Suox*^KO/KO^ mice identified a significant and widespread change in protein persulfidation of more than 400 proteins, most of which were involved in metabolic pathways. Furthermore, consistently with our in vitro studies using Cys-SSH, we found an increase in S-sulfoxidated peptides in liver extracts of *Suox*^KO/KO^ mice. Protein persulfidation protects the respective proteins from sulfonylation (R-SO_3_H) of cysteine residues and a subsequent loss of protein functions. Currently, the effect of reduced protein persulfidation had been mainly studied in the context of aging (Zivanovic et al., 2019), its impacts on the specific proteins remained sparse with only a handful of examples such as the persulfidation on human GAPDH (Cys152), human CSE (Cys252, 255, 307, and 310), and human eNOS (Cys443), which increase the respective enzymes’ activities, while the persulfidation on human PTP1B (Cys215) suppresses its activity (Luo et al., 2023; Vignane & Filipovic, 2023). Therefore, our study identified a large pool of proteins with sulfite-sensitive persulfides that may collectively or individually have contributed to the disease pathology in SOXD. Besides a better understanding of SOXD, we expect that we have identified crucial persulfidation sites in the proteome that may even contribute to stress resistance and longevity.

Sulfite-induced release of H_2_S from persulfides is considered a transient process with yet unknown peak concentration of H_2_S at peak reaction. Given the clinical similarity of MoCD/SOXD with hypoxic ischemia encephalopathy we may speculate that the proposed H_2_S peak would cause inhibition of cytochrome c oxidase and mimic hypoxic conditions. In contrast at moderately increased H_2_S levels, a recent study by the Banerjee laboratory has demonstrated increased mitochondrial respiration due to efficient H_2_S oxidation by SQOR (Landry et al., 2021). When a critical concentration of H_2_S is reached, cytochrome c oxidase will be inhibited, resulting in respiration arrest, increased NADH levels and reversal of the TCA cycle, as demonstrated by glutaminolysis, to replenish α-ketoglutarate. Together with increased glycolysis reducing equivalents will be used for lipogenesis (Carballal et al., 2021). By investigating the amino acid profile in *Suox*^KO/KO^ mice we found a depletion of all catabolic amino acid precursors of α-ketoglutarate, with highest reduction in glutamine levels, thus providing in vivo support for a H_2_S-dependent reversal of the TCA cycle. Given the phenotype difference between MoCD/SOXD and EE patients we propose that beside H_2_S toxicity, additionally, the loss (protein)-persulfidations will additionally contribute to disease severity and will require future studies to dissect the functional impact of respective persulfidations.

## Material and Methods

### CRISPR/Cas9-based generation of *Suox* knockouts in C57BL/6N mice

CRISPR knockouts were generated through a modified protocol from Chu et al. (Chu et al., 2016). Cas9 short guiding (sg)RNA was predicted using Benchling software and selected for highest efficiency and lowest off-target rate on chromosome 10. The selected sgRNA (5’-CCTTACTCATTTCGGAGCGC<u>CGG</u>-3’) was tested on a purified PCR amplicon corresponding to a region in the *Suox* Moco domain in an *in vitro* approach, incubating 30nM amplicon, 30nM sgRNA and 30 nM Cas9 protein (NEB) for 1h at room temperature and analyzing cleavage products on a 2% agarose gel. The main process of generating this model was performed by CECAD research facility, University of Cologne. Fertilized oocytes obtained from breedings of C57BL/6N mice were microinjected with *Suox* gRNA and *Cas9* mRNA and protein. The oocytes were implanted in pseudo-pregnant C57BL/6N mice. Genotypes of off-spring were identified by isolating DNA using QuickExtract buffer (Epicentre, #QE09050), subsequent amplification of the targeted area using primers G1 and G2 and employing the T7 endonuclease assay. In short, the PCR amplicon was isolated and heteroduplices formation was facilitated by incubating the samples at 95°C for 10 min, decreasing the temperature to 85°C at -2°C / min, and finally to 25°C at -0.3°C / min using a PCR cycler. 18 µl of heteroduplexed DNA and 2U T7 endonuclease I (IDT) was incubated at 37°C for 1 h. Cleavage of heteroduplex DNA was analysed on a 2% agarose gel. Target genomic areas of founder mice appearing positive in the T7E1 assay were amplified by PCR, inserted into the pJET vector system (K1232, ThermoFisher Scientific, America) and used for transformation into *E. coli* DH5α; 24 clones were isolated, grown in 5 ml LB medium and plasmid DNA was isolated and subsequently sequenced using Sanger sequencing (Eurofins Genomics) using a T7 promoter primer. Founder mice harboring the desired mutation were bred for two generations before heterozygous matings were started to ensure proper separation of genotypes from chimeric mice.

### Mice-keeping, *Suox* KO mouse breeding and scoring

All animals were kept and bred in accordance with European, national and institutional guidelines and protocols were approved by local government authorities (Landesamt für Natur, Umwelt und Verbraucherschutz Nordrhein-Westfalen, Germany; reference 84-02.04.2014.A372). Mice were kept under a 12 h light cycle and provided with regular chow diet and water *ad libitum*. For the generation of homozygous *Suox* KO mice, heterozygous mice of at least two months of age were kept in 1:1 or 1:2 (male:female) breedings. In case of 1:2 breedings, other females than the mother were removed from the cage before the litters were born. The date of birth (P0) was noted and new litters were observed twice a day until the litter was terminated. Starting at P4, pups were weighed and phenotypically scored.

Animals were sacrificed when weight gain stagnated or animals were found to be suffering. Additionally, righting reflexes of pups were analyzed following a protocol from Didonato et al. Pups were fixated on their backs for approximately five seconds and time to right themselves was recorded. Scores according to righting quality were assigned: 3 – pup rights itself with correct paw placement; 2 – pup rights itself, at least one paw is misplaced; 1 – pup does not right itself, but struggles to for 60 seconds; 0 – pup does not right itself and aborts struggle within 60 seconds. Both male and female mice were analyzed. Genotyping was performed using tail tips and primers after sacrifice. Genomic DNA was extracted using 50µL QuickExtract DNA Extraction Solution (Lucigen, Qiagen). DNA sequences surrounding the deletion site were amplified using the following primers: (1) 5’-TAACACATCACAGAGCCGGG -3’, (2) 5’-CTGGACCCACACACCTATCG -3’. Annealing temperature was 62°C. PCR products were followed incubating at 37°C for 1 hour with restriction enzyme, Bfol (FastDigest Bfol, FD2184, ThermoFisher Scientific, America). The entire products were run in 2% TBE gel for 30 mins at 100V.

### Western blot analysis

Western blotting was performed on crude protein extracts from different tissues lysed in 100 mM Tris/Ac, pH 8.0. Tissue samples were weighed before lysis and lysis buffer volume was adjusted accordingly. Unless otherwise indicated, 15 µg of protein lysate were separated by SDS-PAGE and immunoblotted using standard protocols. Membranes were probed using HRP-coupled secondary antibodies; signals were detected using chemiluminescent substrates (ThermoFisher Scientific, #34580) and a BioRad ChemiDoc XRS+ system. For densitometric measurements, images were converted to 8-bit using ImageJ (NIH, version 1.50i). Lanes were selected using the square tool and histograms for individual bands were evaluated after background removal. Identical squares were used for each quantified band. Unless otherwise stated, signals were normalized to VCL or GAPDH as loading control.

### SOX activity measurements

SOX activity was measured using the sulfite:cytochrome c activity assay in multiple tissues as described previously (Kumar et al., 2017). In short, tissue samples were removed from animals after sacrifice, flash-frozen in liquid nitrogen and stored -80°C until further usage. Tissues were lysed in 100 mM Tris/Ac, pH 8.0 using a loose-fitting homogenizer at 1200 rpm for 10 cycles, and subsequent sonication for 60s at 15% using 3s / 3s cycles. Lysates were centrifuged at 21.000 rcf for 45 min at 4°C. Protein concentration was determined using the Bradford assay. In general, 20 – 200 µg crude protein extract was incubated in 200 µl 50 mM Tris/Ac, pH 8.0, 0.2 mM deoxycholic acid, 0.1 mM potassium cyanide and 0.5 mM sulfite. The reaction was started by the addition of 100 µl of 100 mM equine cytochrome *C*. SOX activity was determined by monitoring the absorption change at 550 nm (ε_550_ = 19630 M^-1^ cm^-1^) using a 96 well-plate-based approach and a BioTeK plate reader at room temperature.

### Immunohistochemistry

*Suox^WT/WT^* and *Suox^KO/KO^* pups (P8) of either sex were anesthetized with 100 mg/kg xylazine (Rompun 2%, Bayer) and 20 mg/kg ketamine (Ketaset, zoetis) and perfused first with PBS and then 4 % (w/v) formaldehyde in PBS. Brains were post fixed for 24 h in 4 % formaldehyde/PBS. Afterwards brains were washed with 50 mM NH_4_Cl in PBS and cryoprotected first in 15 and then 30 % (w/v) sucrose in PBS. Free-floating coronal slices of the right hemisphere (30 µm thickness) were prepared on a Leica CM3050S cryostat at -16°C and placed in PBS at 4°C. All subsequent incubation and wash steps were performed at RT.

After blocking/permeabilization for 1 h with 10 % goat serum, 0.4 % TritonX-100 in TBS, the following primary antibodies were used: anti-NeuN specific (1:500, ThermoFisher # PA5-78639) and anti-MAP2 specific (1:1000, Millipore #AB5622). The following secondary antibody was used: Goat anti-rabbit AlexaFluor 647 (1:500, #A-21245, Thermo Fisher Scientific). Nuclei were stained with 3 µM DAPI in PBS for 5 min. at RT. Slices were mounted with Mowiol/Dabco.

### Confocal microscopy and image analysis

Images were acquired on a Leica TCS SP8 LIGHTNING upright confocal microscope, equipped with hybrid detectors (Leica HyD) and the following diode lasers: 405 nm, 488 nm, 552 nm, and 638 nm. Image tiles of entire slices (per tile: 1,024 x 1,024 (1163.34 µm x 1163,64 µm)) were recorded with an HC PL APO CS2 10x/0.40 dry objective and high magnification images (3,432 x 3,432 (144.77 µm x 144.77 µm)) with an HC PL APO CS2 63x/1.30 Glyc objective. LIGHTNING adaptive deconvolution using ‘Mowiol’ setting were applied on high magnification images. Images were segmented and analyzed in an automated fashion using ImageJ/FIJI 1.53c (https://imagej.net/software/fiji/). Rolling ball background subtraction, median filtering, and Otsu auto thresholding methods were used for image segmentation.

### Statistical analysis and data visualization (IHC)

Individual data points per mouse (5 animals per genotype) and mean are displayed in the figures. Data were analyzed with unpaired Student’s t-test. Statistical analyses and data visualization were performed using R version 4.1.0 (2021-05-18), rstatix v. 0.7.0, RcolorBrewer v. 1.1-2, dplyr v. 1.0.7, purrr v. 0.3.4, readr v. 2.0.0, tidyr v. 1.1.3, tibble v. 3.1.3, ggplot2v. 3.3.5, tidyverse v. 1.3.1, rio v. 0.5.27, pacman v. 0.5.1 in Rstudio environment (https://www.rstudio.com/), Platform: x86 64-w64-mingw32/x64 (64-bit). Figures were prepared with: OMERO (https://www.openmicroscopy.org/).

### Sulfite and thiosulfate determination in plasma and urine

Plasma was collected by mixing whole blood with EDTA and centrifugation. Both plasma and urine were stored at -80 °C until further measurement. Sulfite and thiosulfate were determined using monobromobimane derivatization and subsequent HPLC analysis (Hildebrandt et al., 2013). 15 µl of sample were mixed with 15 µl 160 mM HEPES, pH 8.0, 16 mM EDTA, 15 µl acetonitrile and 3 µl 46 mM monobromobimane in acetonitrile. The samples were mixed and incubated in the dark for 30 min at room temperature. The reactions were stopped with either 65 mM methanesulfonic acid (for urine) or 1.5% methanesulfonic acid (for plasma). The samples were diluted with a four-fold volume of solvent A (0.5% acetic acid, pH 5.0). Flow rate was set to 1 ml / min. 5 or 80 µl of sample mix were injected into a LiChrospher 60 RP-Select B (5 µm) 125-4 column. The program is described with % buffer B (100% MeOH): 0.0 min, 8.0%; 7.5 min, 8.0%; 7.51 min, 12.0%; 17.5 min, 12.0%; 17.51 min, 100.0%; 19.0 min, 100%; 19.01 min, 8.0%; 25.0 min, 8.0%. The sulfite-bimane peak eluted at 3.4 min, the thiosulfate-bimane peak at 6.8 min with fluorescent detection at 380nm excitation and emission at 480nm.

### *S*-sulfocysteine determination in plasma and urine

*S*-sulfocysteine was determined as described previously (Jakubiczka-Smorag et al., 2016). Protein in plasma samples was precipitated through addition of two volumes of ice-cold methanol. Samples were incubated on ice for 15 min and subsequently centrifuged at 21.000 g for 10 min at 4°C. Urine samples were centrifuged at 21.000 g for 10 min at 4°C. 15 µl of supernatant from either preparation were mixed with 15 µl 200 mM Na_2_CO_3_/NaHCO_3_, pH 9.5 and 30 µl 20 mM 9-fluorenylmethyloxycarbonyl chloride (FMOC-Cl). The reaction was incubated at room temperature for 20 min. The reaction was stopped by the addition of 5 µ 80 mM amantadine hydrochloride (dissolved in 1:1 acetonitrile / 0.2 M HCl). After 5 min incubation time, the sample was centrifuged at 21.000 g and the supernatant was transferred into a glass HPLC vial. SSC was separated on a 250 x 3 mm 5 µm Nucleodur HTec C18 (Macherey-Nagel) column. Solvent (A) was 50 mM Sodium formate, 0.05% trifluoroacetic acid, pH 4.5 in water; solvent (B) was 100% methanol. Flow rate was set to 1 ml / min; a gradient from 35 to 60% B was performed from 0 to 7.5 min; SSC eluted at 8.81 min SSC peak area was quantified at 265 nm.

### Amino acid quantification in mouse plasma and urine

Amino acids in mouse plasma and urine were measured by automated column amino acid analyser (Biochrom 30), according to standardized procedures (Duran, 2008).

### MRI on mouse brains

A 3.0 T Philips Achieva clinical MRI scanner (Philips Best, the Netherlands) in combination with a dedicated mouse solenoid coil (Philips Hamburg, Germany) was used for imaging. Three dimensional T2-weighted MR images were acquired post mortem, using a turbo-spin echo (TSE) sequence with repetition time (TR) = 439 ms, echo time (TE) = 48 ms, field of view (FOV) = 60 x 35 x 25 mm^3^, voxel size = 0.2 x 0.2 x 0.2 mm^3^, number of averages = 3. MR images were analyzed with the software VINCI 4.72 (Max Planck Institute for Metabolism Research, Cologne, Germany). Brain volume was determined by manually fitting a volume of interest (VOI) to each individual brain.

### *In vitro* and monobromobimane-based H_2_S detection

NAP-SSH was synthesized as described by Artaud and Galardon, 2014 (Artaud & Galardon, 2014). TNB-SSH was generated by adding an equimolar amount of NaHS to DTNB in 50 mM PBS, pH 7.4. HSA-SSH and GAPDH-SSH were generated as described previously (Cuevasanta et al., 2015). For the detection of H_2_S release from persulfidated compounds a selective H_2_S electrode of a Free Radical Analyzer (World Precision Instruments) was used. For detection of bioavailable H_2_S in plasma, monobromobimane labeling followed by HPLC with fluorescent detection was employed based on a standardized method. (PMID: 30970274). 25 μl of the plasma sample was mixed with 66 μl of freshly mixed working buffer (65 μl 200mM HEPES pH 8.2 + 100mM monobromobimane in ACN) in the dark at precisely 20.0 °C for exactly 10 minutes. The reaction was stopped by the addition of 5 μl 50% TCA, precipitated proteins were removed by centrifugation at 3000g for 5 minutes RT and the supernatants were transferred into amber HPLC autosampler vials. 2 μl was injected onto a Phenomenex Luna C18(2) column (250×2 mm, 3μm) and separated by the following optimized gradient elution profile using 0.1%TFA/H_2_O (A) and 0.1%TFA/ACN (B) with 0.2 ml/min flow rate: The gradient started with 15%B increasing to 35% in 3.54 min and kept there for 7.09 min. In 2.36 min the composition was increased to 90%B, held for 1.18 min and returned to the initial 15% in 1.18 min. A Thermo Scientific Ultimate 3000 HPLC system was used for the separation, for the fluorescent detection an excitation wavelength of 390 nm and an emission wavelength of 475 nm was employed.

### Detection of persulfidated proteins in tissue

#### Proteomic analysis of the persulfidome

Persulfidation labeling was performed as described previously (Zivanovic et al., 2019). Briefly, liver samples were lysed in cold HEN Buffer (50 mM HEPES, 1 mM EDTA, 0.1 mM Neocuproine, 1 % IGEPAL, 2 % SDS, pH 7.4) supplemented with 10 mM 4-Chloro-7-nitrobenzofurazan (NBF-Cl, Sigma, #163260) and 1% Protease Inhibitor using TissueLyser II (Qiagen) and 5 mm stainless steel beads (Qiagen, #69989). Lysates were incubated for 2 h at 37°C protected from light. Methanol/chloroform precipitation was performed twice (MeOH/CHCl3/sample, 4/1/1, v/v/v) and pellets washed with cold methanol. After resuspension in 2% SDS in PBS (Sigma-Aldrich, #P4474) samples were cleaned from endogenously biotinylated proteins using Pierce™ NeutrAvidin™ Agarose beads (Thermofisher, #29200) and precipitated. Pellets were then resuspended in PBS, 2% SDS, incubated for 1.5 h at 37°C with 250 µM DCP-Bio1 (MERK, #NS1226) and precipitated again. Protein concentration was adjusted to the same level using DC protein assay. After 2 h incubation on Ferris wheel at room temperature (RT) and protected from light, supernatants were collected and precipitated. Proteins were redissolved and their concentration was adjusted to the same level using DC protein assay. Equal amount of proteins was mixed with Pierce™ High Capacity Streptavidin Agarose beads and incubated at RT for 4 h in a Ferris wheel protected from light. The samples were transferred into 2 mL Pierce™ Disposable Columns (Thermofisher, #29920) where washing step was carried out with 28 ml of PBS and 8 ml of water to remove nonspecifically bound proteins and detergent. Elution step of enriched proteins was performed using 2.25 M ammonia solution (Sigma, #5.33003) for 16 h. Samples were lyophilized and dissolved in digestion buffer (50 mM ammonium bicarbonate [ABC], 1 mM calcium chloride) and digested overnight using trypsin (Promega, #V5117) at trypsin: protein ratio 1:20. The desalting step was performed on Supel™-Select HLB SPE Tube (Sigma, #54181-U).

Peptides were dissolved in 0.1 % TFA, and digestion quality control was performed using an Ultimate 3000 Nano Ultra High-Pressure Chromatography (UPLC) system with a PepSwift Monolithic® Trap 200 µm * 5 mm (Thermo Fisher Scientific). Peptides were analyzed by high-resolution LC-MS/MS using an Ultimate 3000 Nano Ultra High-Pressure Chromatography (UPLC) system (Thermo Fisher Scientific) coupled with a Thermo Orbitrap Eclipse™ Tribrid™ Mass Spectrometer via an EASY-spray (Thermo Fisher Scientific). Peptide separation was carried out with an Acclaim™ PepMap™ 100 C18 column (Thermo Fisher Scientific) using a 120 min non-linear gradient from 3 to 35 % B (0 min, 3 % B;5 min, 7 % B; 120 min 35 % B; A: H_2_O, B: 84 % Acetonitrile, 0.1 % Formic Acid) at a flow rate of 250 nL/min. The Orbitrap Eclipse was operated in a DDA mode, and MS1 survey scans were acquired from 300 to 1500 m/z at a resolution of 120,000 using the Orbitrap mode. The ten most intense peaks with a charge state ≥ 2 were fragmented using Higher-energy collisional dissociation (HCD) at 32 % and analyzed using Ion-trap at a normal scan rate. The dynamic exclusion was set at 30 seconds.

Data were evaluated with PEAKS ONLINE software using 15 ppm for precursor mass tolerance, 0.5 Da for fragment mass tolerance, specific tryptic digest, and a maximum of 3 missed cleavages. NBF (+163.00961594 Da) on C, K, R, DCP (+168.0786442 Da) on C, N-term acetylation (+42.010565 Da), and methionine oxidation (+15.994915 Da) were added as variable modifications, peptide-spectrum match (PSM) and proteins were filtered at FDR 1 %. Data were normalized using the eigenMS (Karpievitch et al., 2009) script on R studio.

#### Total proteome analysis

For the total proteome analysis, 100 µg of proteins obtained by the above-mentioned lysis step, were digested overnight at 37 °C using trypsin with a 1:20 trypsin to sample ratio (Promega, #V5117). The next day, peptides were desalted and evaporated under vacuum until dryness as described above.

Peptides were dissolved in 0.1 % TFA, and digestion quality control was performed as described above. Peptides were analyzed by high-resolution LC-MS/MS using an Ultimate 3000 Nano Ultra High-Pressure Chromatography (UPLC) system coupled with a Thermo Orbitrap Eclipse™ Tribrid™ Mass Spectrometer via an EASY-spray. Peptide separation was carried out with an Acclaim™ PepMap™ 100 C18 column using a 150 min linear gradient from 3 to 35 % of B (0 min, 3 % B; 140 min, 28 % B; 150 min 35 % B; A: H_2_O, B: 84 % Acetonitrile, 0.1 % Formic Acid) at a flow rate of 250 nL/min. Thermo Orbitrap Eclipse™ was operated in DDA mode using Thermo Xcalibur Instrument Setup Software. For the MS scan the detector was operated in Orbitrap with a scan range between 300 and 1500 with positive polarity including charge states between 2 and 7. For the MS2 we used a Quadrupole isolation mode with a HCD activation type and the dynamic exclusion parameter were of 30 seconds for the exclusion duration.

Data were evaluated with PEAKS ONLINE software using 15 ppm for precursor mass tolerance, 0.5 Da for fragment mass tolerance, specific tryptic digest, and a maximum of 3 missed cleavages. Carbamidomethylation (+57.021464 Da) on C, N-term acetylation (+42.010565 Da), and methionine oxidation (+15.994915 Da) and NBF (+163.00961594 Da) on C, K, R, were added as variable modifications, PSM and proteins were filtered at FDR 1 %. Data were normalized based on the total ion count (TIC).

### Proteomic analysis of tissues

Whole tissues were excised from mice after decapitation, washed with PBS and flash-frozen until processing. For homogenization, tissues were grounded in a liquid nitrogen-cooled mortar to fine powder. The tissue powder was suspended in 50 mM triethylammoniumbicarbonate, 8 M urea, supplemented with protease inhibitor (cOmplete EDTA-free, Roche) and phosphatase inhibitor (PhosSTOP, Roche). Chromatin was degraded using a Bioruptor sonication rod (Diagenode Diagnostics) for 10 min using 30s-30s pulse-break cycles on ice. After centrifugation and protein concentration determination, 50 µg protein were transferred to a new Eppendorf tube and reduced via addition of DTT (5 mM, 37°C, 1 h). Alkylation was performed by addition of chloroacetamide (40 mM, 37°C, 2 h). Proteolytic cleavage was performed by an initial addition of lysylendopeptidase C (37°C, 4 h) and subsequent addition of trypsin (37°C, overnight). Peptides were purified by mixed chromatography using styrenedivinylbenzene-RP. In short, two disks of styrenedivinylbenzene-RP-sulfonate were stacked in pipette tips (StageTips) and equilibrated by consecutive application of 1) methanol, 2) 0.1% formic acid in 80 % acetonitrile, 3) twice 0.1 % formic acid. The sample was applied and washed multiple times using 1) 0.1 % formic acid and 2) thrice 0.1 % formic acid in 80 % acetonitrile. The StageTips were dried and stored at 4°C until further processing. Analysis was conducted by CECAD Proteomics Core Facility. In short, peptides were separated using C18-RP chromatography (Poroshell EC120, Agilent; 50 cm length, 75 µm diameter, 2.7 µm particle size) on an EASY nLC (Thermo Scientific). Peptides were loaded with solvent A (0.1 % formic acid) and separated at 250 nl min^-1^ in a gradient: 3 – 4 % solvent B (0.1 % formic acid in 80 % acetonitrile) within 1.0 min, 4 – 27 % solvent B within 119.0 min, 27 – 50 % solvent B within 19.0 min, 50 – 95 % solvent B within 1.0 min. The mass spectrometer was operated in data-dependent acquisition mode.

## Data and materials availability

All proteomic data are available through PRIDE under code number PXD045896 (**Project DOI:** 10.6019/PXD045896, **Username:**, **Password:** KH0K3hbQ).

## Supporting information

Fu_Kohl_etal_Suppl

## Acknowledgements

GS and CYF were supported by the German Research Foundation (Deutsche Forschungsgemeinschaft, DFG) SFB1218 project number 269925409 and SFB1403 project number 414786233. MRF acknowledges the support of European Research Council (ERC) under the European Union’s Horizon 2020 research and innovation programme (Grant Agreement No. 864921). PN and TD acknowledges support from the National Research, Development and Innovation Fund of the Ministry of Culture and Innovation under the National Laboratories Program (National Tumor Biology Laboratory (2022-2.1.1-NL-2022-00010)) and the Hungarian Thematic Excellence Program (under project TKP2021-EGA-44) Grant Agreements with the National Research, Development and Innovation Office. PN acknowledges the support of the Hungarian Research Network – HUN-REN–UVMB Laboratory of Redox Biology Research Group (grant 15002). VK and MK were supported by the grant NU23-07-00383 (Czech Health Research Council), and by institutional programs 21 RVO-VFN 64165 (General University Hospital in Prague) and Cooperatio-Metabolic Disorders (Charles University).

## Supplemental figures

**Supplemental figure 1.**
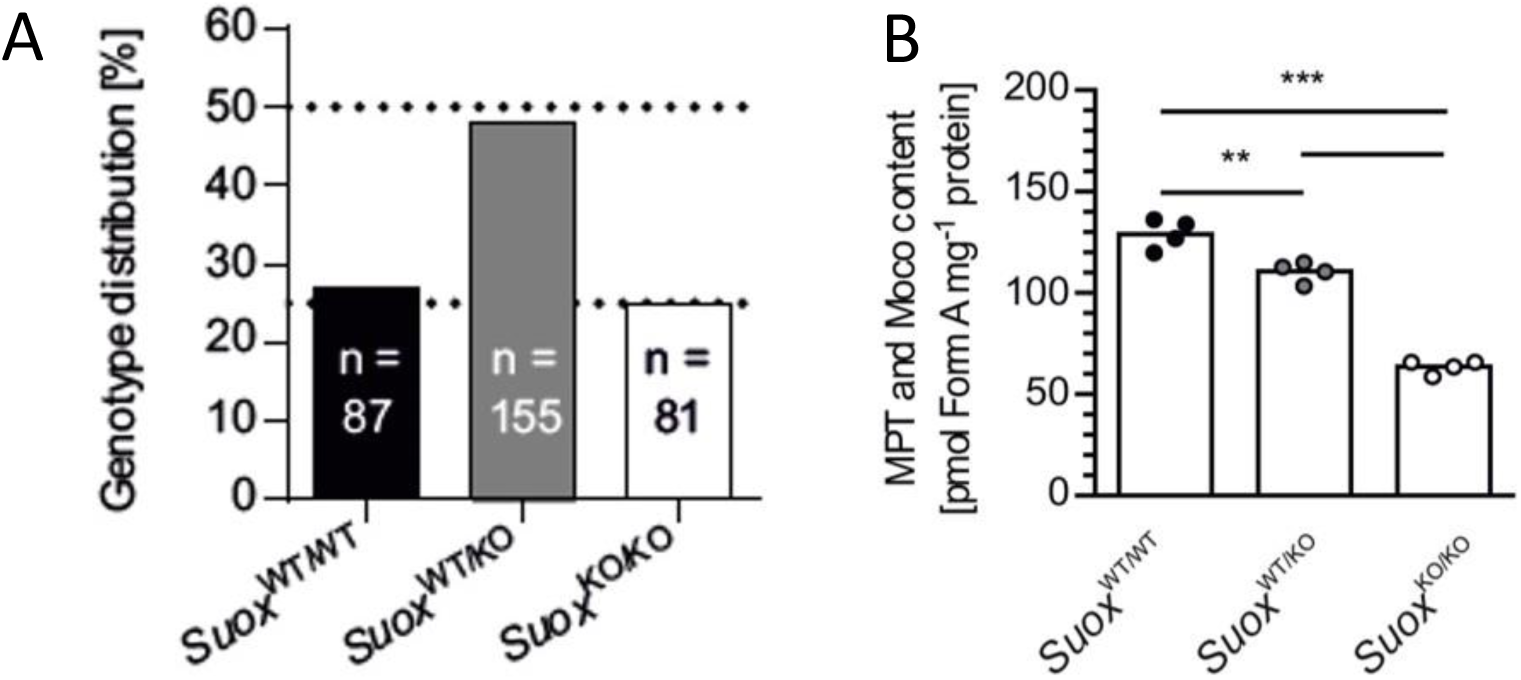
Fu, Kohl et al.

**Supplemental figure 2.**
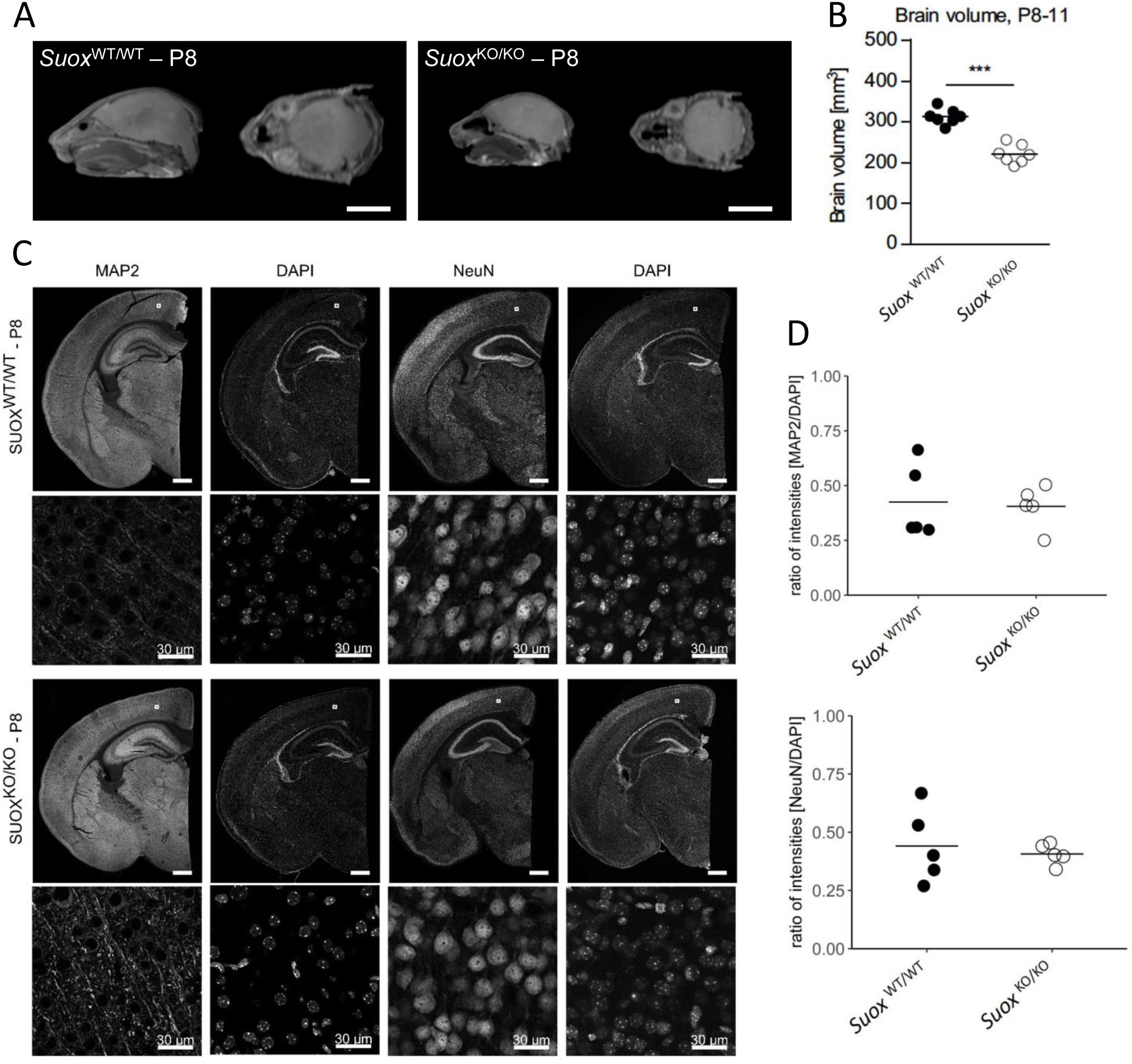
Fu, Kohl et al.

**Supplemental figure 3.**
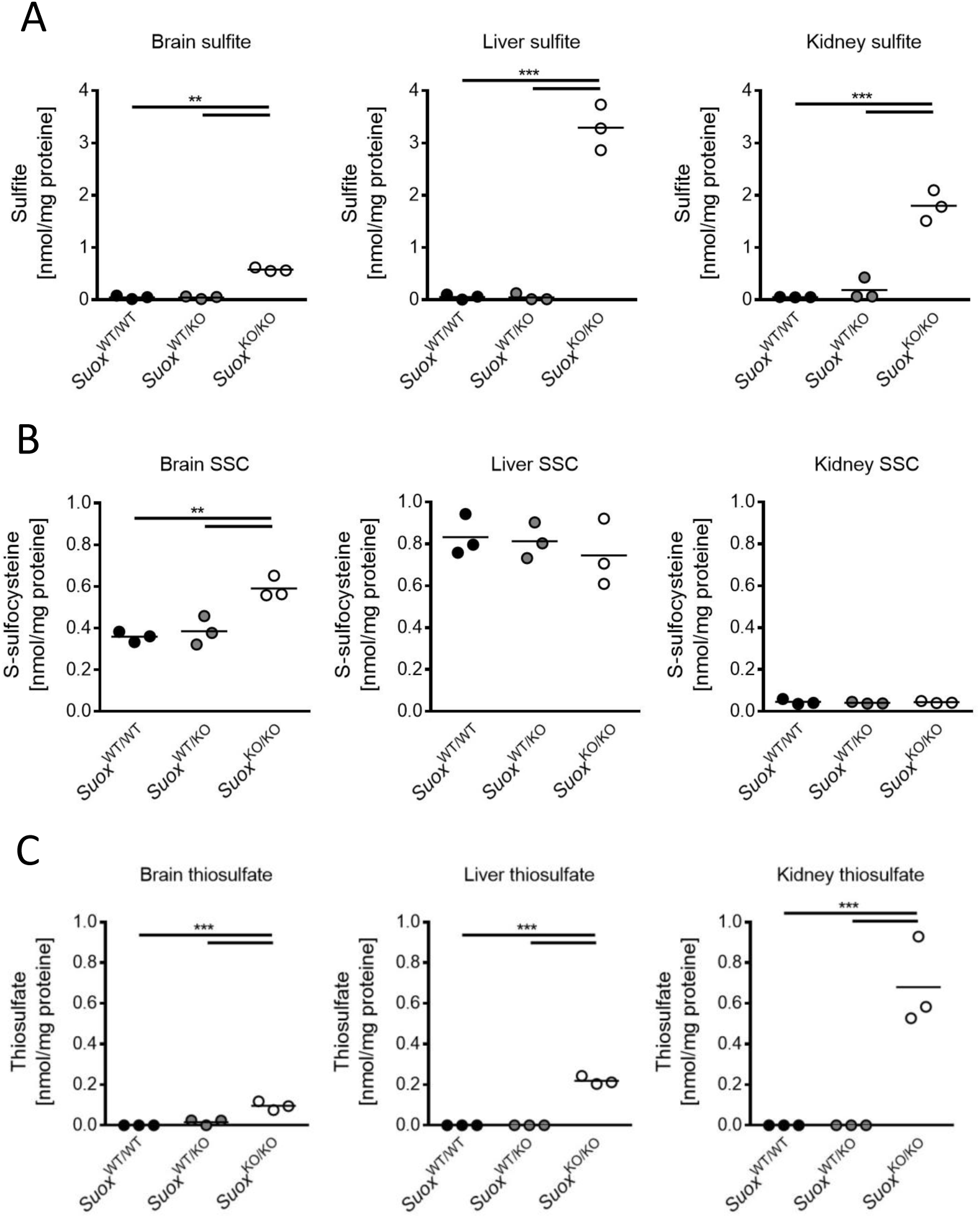
Fu, Kohl et al.

**Supplemental figure 4.**
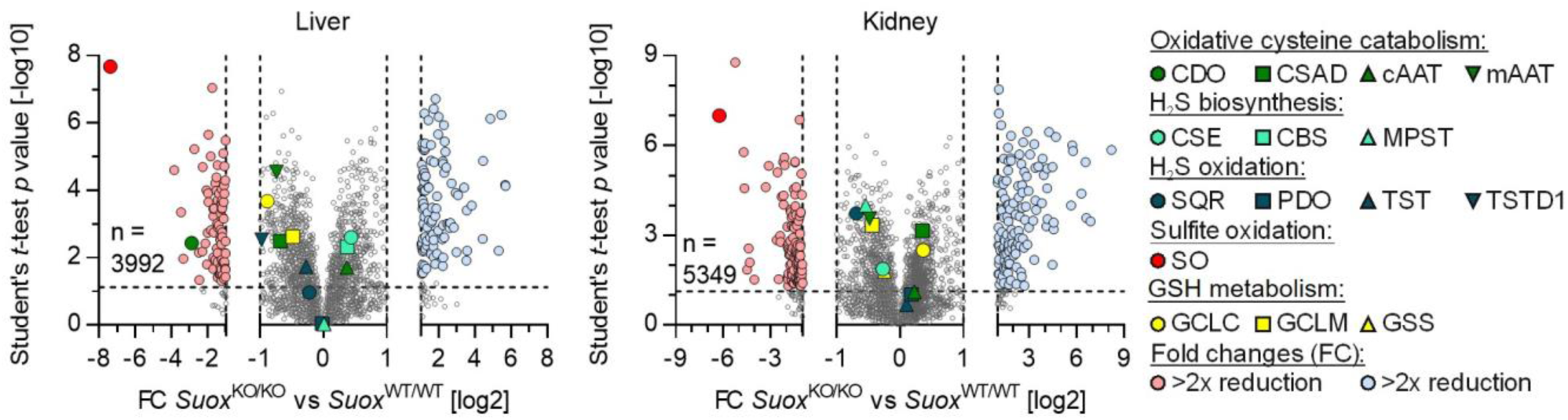
Fu, Kohl et al.

**Supplemental figure 5.**
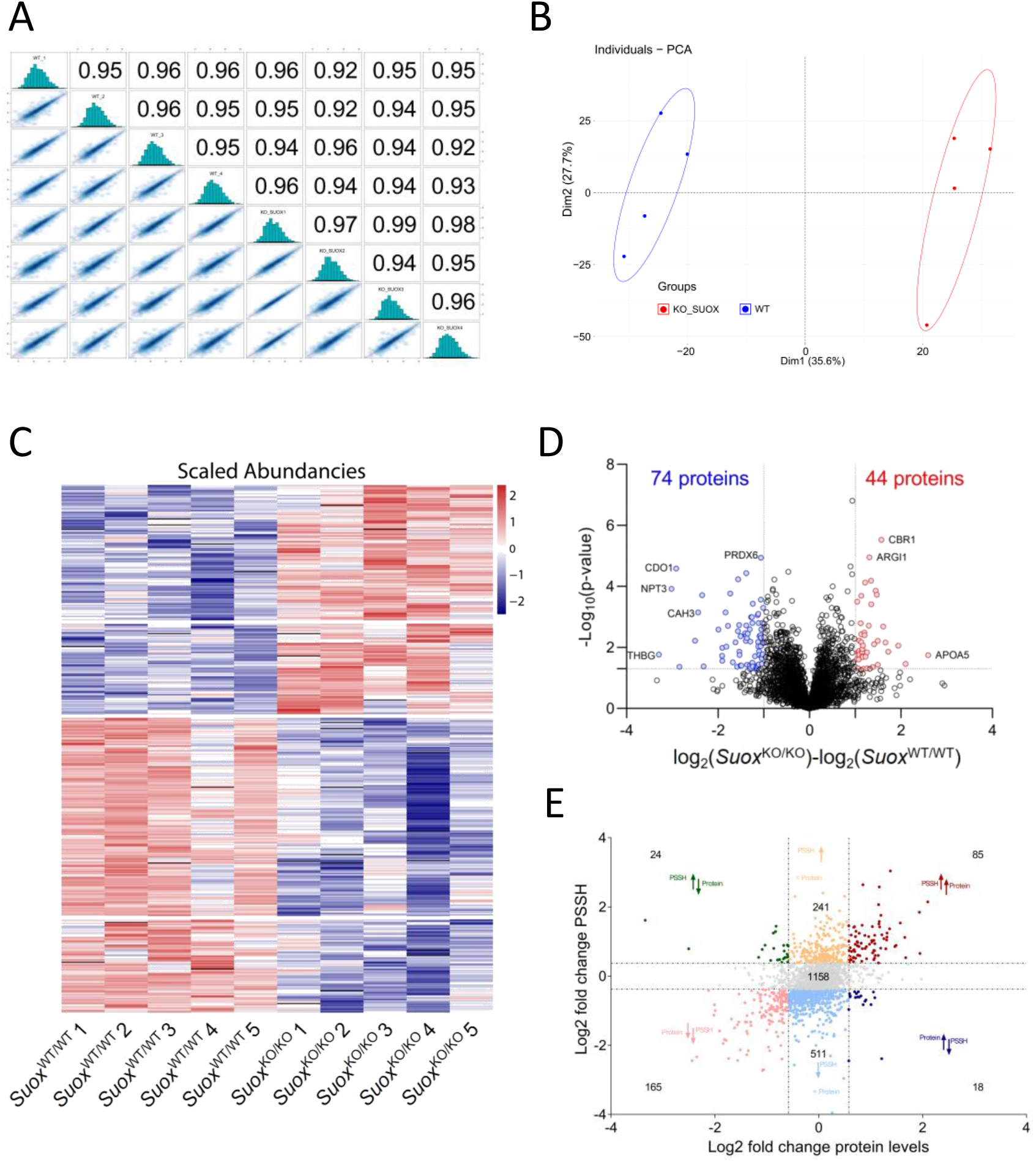
Fu, Kohl et al.

**Supplemental figure 6.**
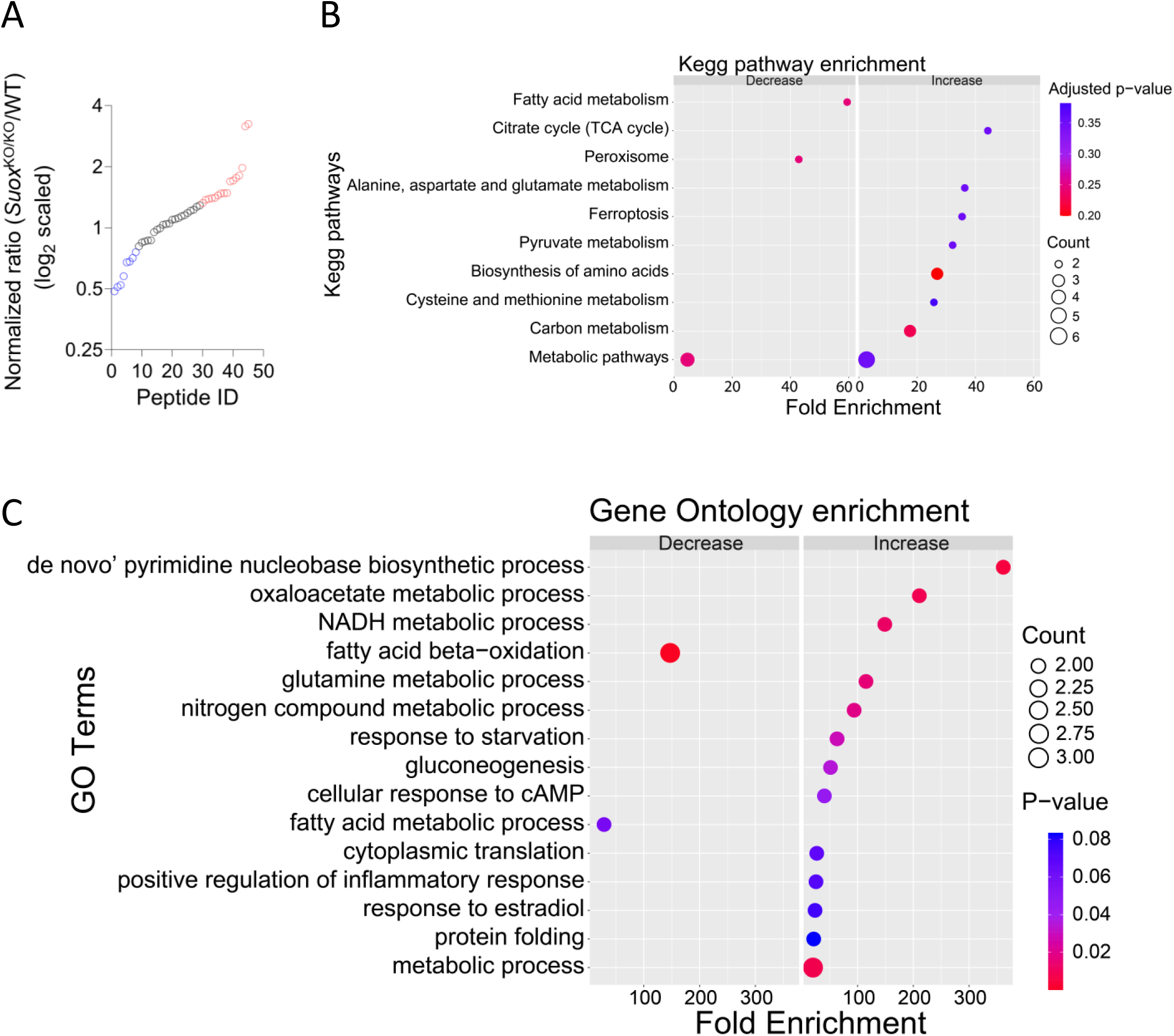
Fu, Kohl et al.

