## Supplementary material for "Sulfite oxidase deficiency causes persulfidation loss and H_2_S release": Fu_Kohl_etal_Suppl

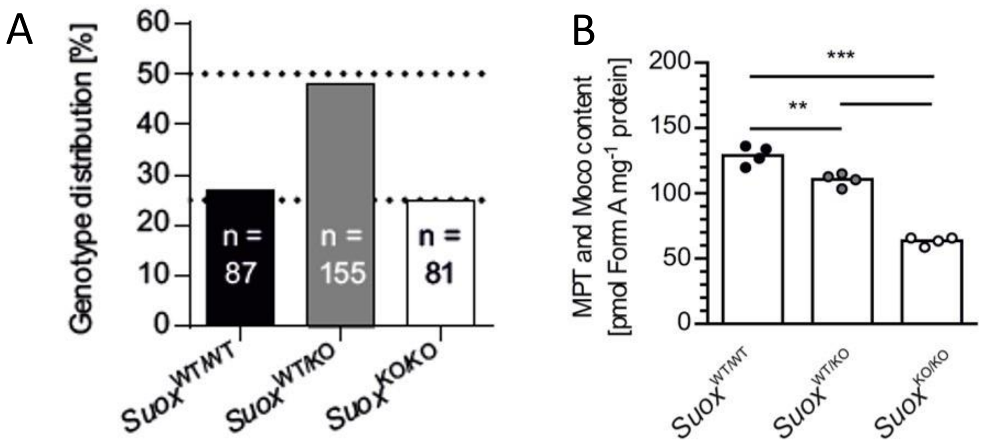


**Supplemental Figure 1. Characterization of *Suox*$^{KO/KO}$ mice.**

(A) Genotype distribution in offspring from *Suox*$^{WT/KO}$ x *Suox*$^{WT/KO}$ breedings. Dotted lines indicate Mendelian distribution. (B) Determination of MPT/Moco content by HPLC FormA analysis in *Suox*$^{WT/WT}$, *Suox*$^{WT/KO}$, and *Suox*$^{KO/KO}$ mice.

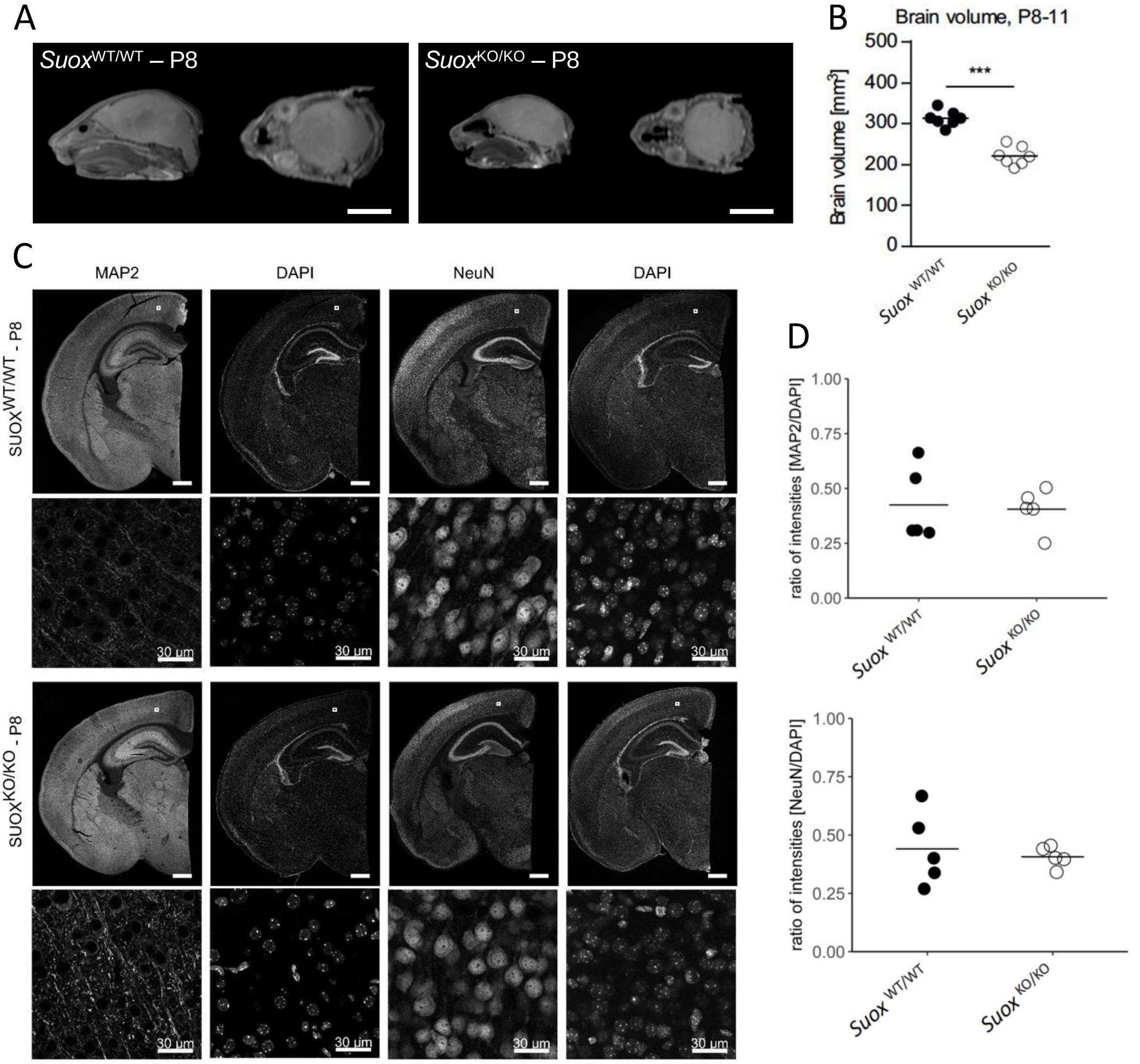


**Supplemental Figure 2. Decreased brain volume in *Suox*$^{KO/KO}$ mice, yet no apoptotic neurons were observed.**

**(A)** Brain magnetic resonance imaging (MRI) scans of P8 *Suox*$^{WT/WT}$ and *Suox*$^{KO/KO}$ mice. Scale bar: 5 mm. **(B)** Decreased brain volume of SOX-deficient mice (P8–P11, n = 7 per group, groups are exactly age-matched), p<0.001 (Student's t test). **(C)** IHC staining for microtubule-associated protein 2 (MAP-2) and neuronal nuclear antigen (NeuN). **(D)** Neuron density showed no difference between *Suox*$^{KO/KO}$ and WT mice.

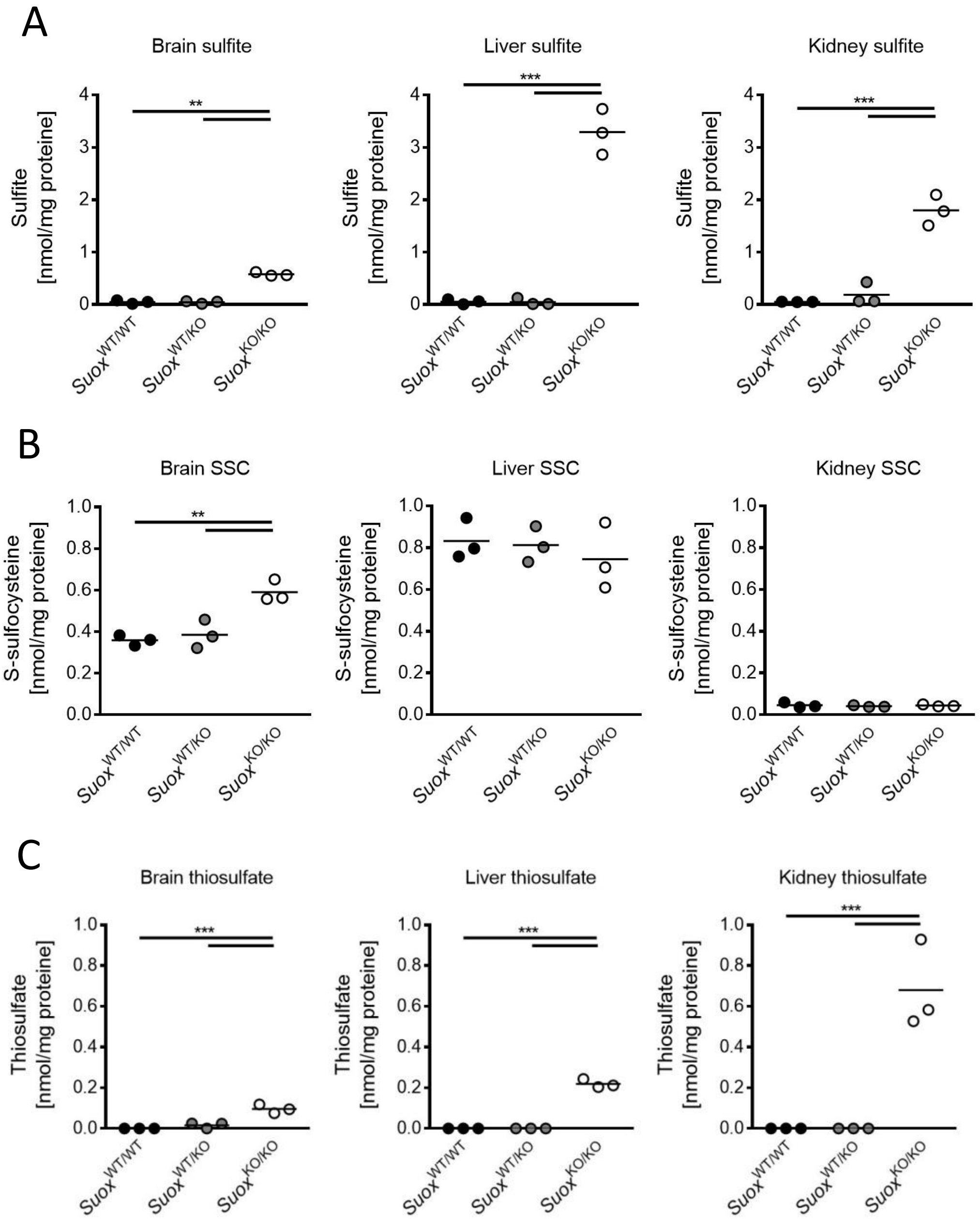


**Supplemental Figure 3. Major SOXD biomarkers in tissues in *Suox*$^{KO/KO}$ mice.**

**(A)** Determination of sulfite, **(B)** S-sulfocysteine, and **(C)** thiosulfate in brain, liver and kidney of *Suox*$^{WT/WT}$, *Suox*$^{WT/KO}$, and *Suox*$^{KO/KO}$ mice. One-way ANOVA with Tukey's post-hoc test for pairwise comparisons was performed as indicated. $p$ value: *** < 0.001; ** < 0.01; * < 0.05; ns > 0.05.

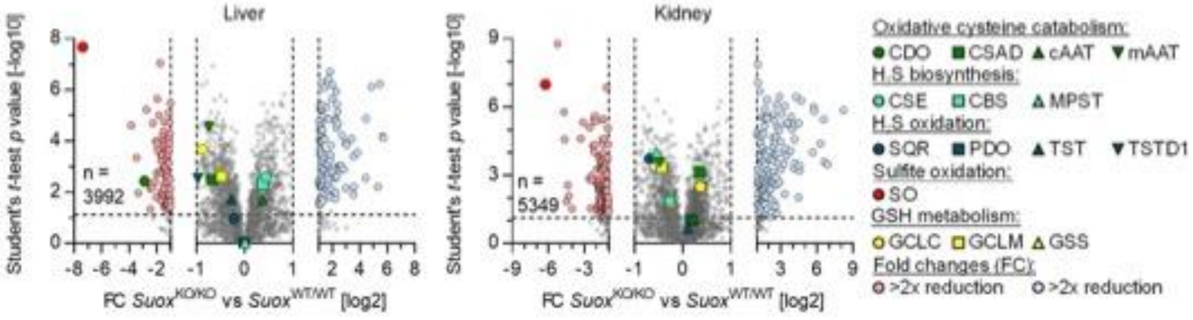


**Supplemental Figure 4. Proteomic profiling of of *Suox*$^{KO/KO}$ mice liver and kidney extracts.**

Volcano plot depicting significantly regulated proteins of *Suox*$^{KO/KO}$ livers compared to *Suox*$^{WT/WT}$ samples (n = 5). Significantly up-regulated proteins are displayed in blue, significantly down-regulated proteins are displayed in red (absolute log2 fold change ≥ 1, -log10 *p* value ≥ 1.3). Exemplary proteins are labelled with their respective gene names.

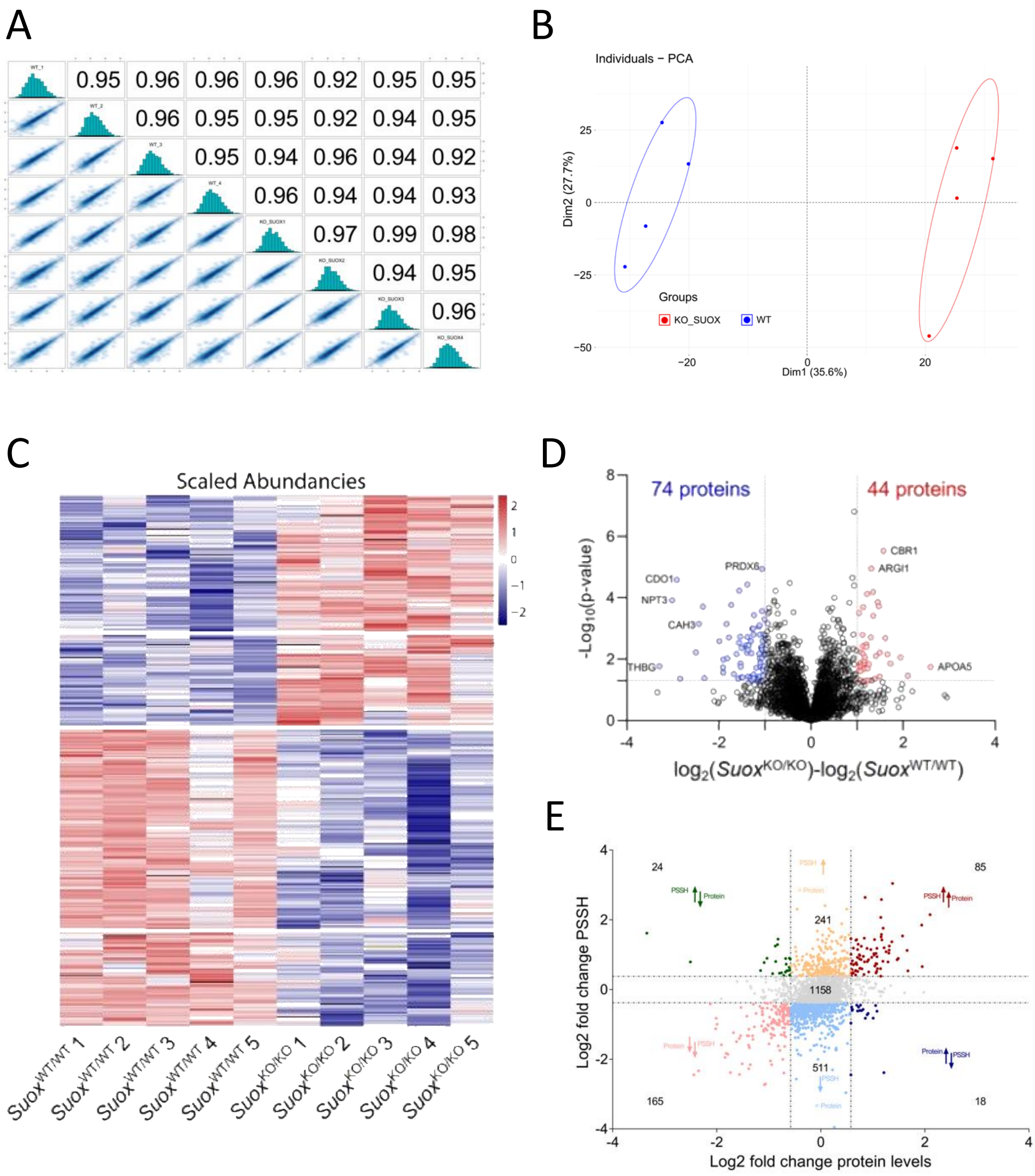


**Supplemental Figure 5. Changes in total proteome of *Suox*$^{KO/KO}$ mouse liver extracts.**

**(A)** Peptide correlation between all measured samples for persulfidome analysis. **(B)** Principal Component Analysis (PCA) shows notable difference in persulfidome of *Suox*$^{KO/KO}$ vs *Suox*$^{WT/WT}$ mice. **(C)** Heatmap showing the changes of protein expression (total proteome) in *Suox*$^{KO/KO}$ mouse livers compared to *Suox*$^{WT/WT}$ mouse (Welch's test, $p<0.05$). **(D)** Volcano plot depicting statistical significance plotted against the $log_2$-fold change of protein expression (total proteome) in *Suox*$^{KO/KO}$ mouse liver relative to *Suox*$^{WT/WT}$ mouse. Significance was established using Welch's t-test (two-sided), with a p-value threshold of < 0.05. Fold change cut-offs were established at 50%. **(E)** Scatter plot analysis of persulfidome and protein expression changes correlation. Scatter plot of the log2 folds changes within the persulfidome (y-axis) and the protein expression (x-axis) for the comparison between *Suox*$^{KO/KO}$ mice and *Suox*$^{WT/WT}$ mice.

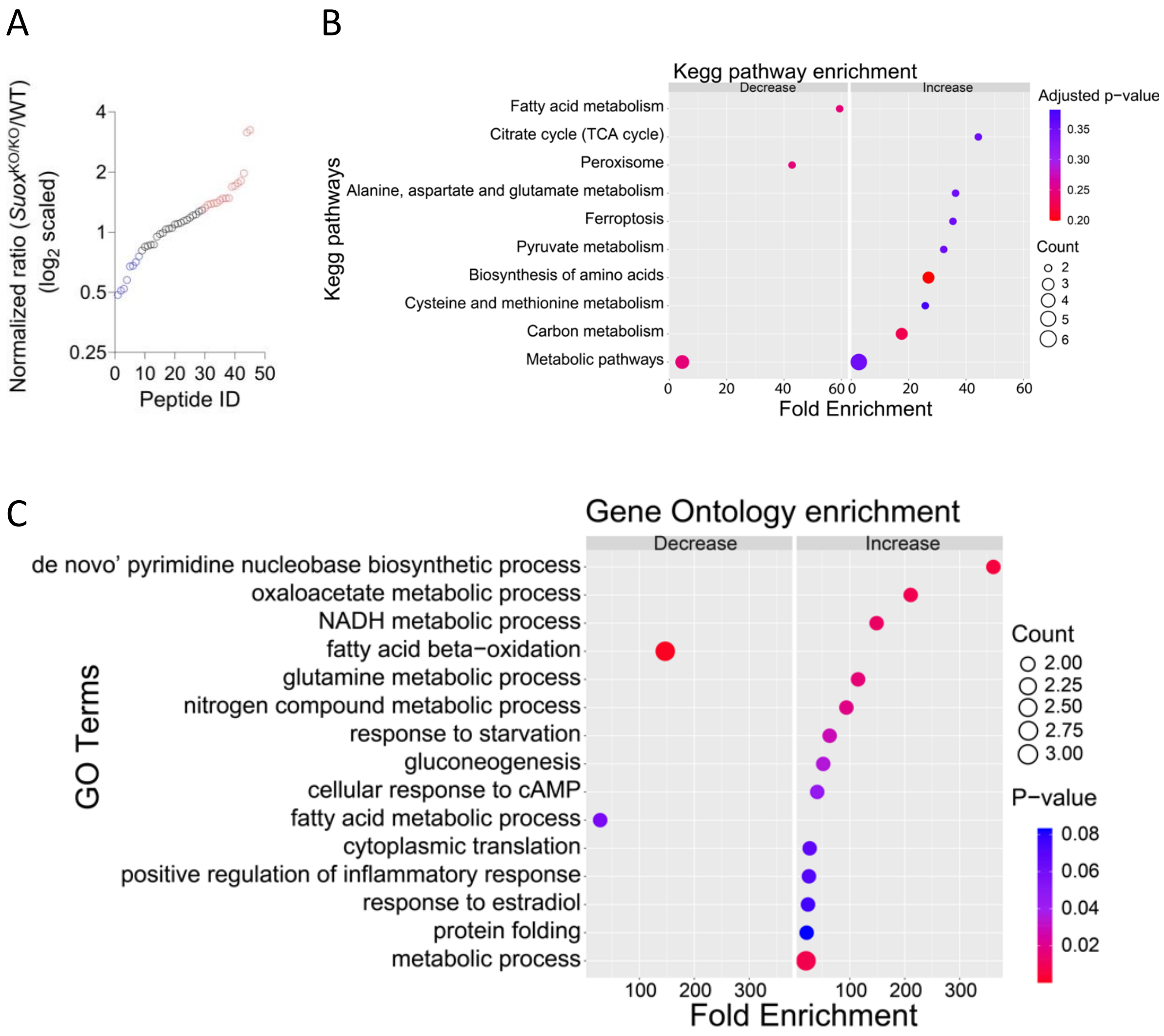


**Supplemental Figure 6. Proteome analysis of peptides with S-sulfonylated cysteines within the total proteome of *Suox*$^{KO/KO}$ mouse liver extracts.**

**(A)** PSSO$_3$H fold change levels normalized to the corresponding protein expression levels.

**(B-C)** Kegg pathway (**B**) and GO Term (biological process) (**C**) enrichment analysis of proteins found to significantly decrease or increase their PSSO$_3$H levels in *Suox*$^{KO/KO}$ mouse liver extracts.
